# CADET: Enhanced transcriptome-wide association analyses in admixed samples using eQTL summary data

**DOI:** 10.1101/2024.10.21.619441

**Authors:** S. Taylor Head, Qile Dai, Joellen Schildkraut, David J. Cutler, Jingjing Yang, Michael P. Epstein

## Abstract

A transcriptome-wide association study (TWAS) is a popular statistical method for identifying genes whose genetically-regulated expression (GReX) component is associated with a trait of interest. Most TWAS approaches fundamentally assume that the training dataset (used to fit the gene expression prediction model) and testing GWAS dataset are from the same ancestrally homogenous population. If this assumption is violated, studies have shown a marked negative impact on expression prediction accuracy as well as reduced power of the downstream genetrait association test. These issues pose a particular problem for admixed individuals, whose genomes represent a mosaic of multiple continental ancestral segments. To resolve these issues, we present CADET, which enables powerful TWAS of admixed cohorts leveraging the local-ancestry (LA) information of the cohort along with summary-level eQTL data from reference panels of different ancestral groups. CADET combines multiple polygenic risk score models based on the summary-level eQTL reference data to predict LA-aware GReX components in admixed test samples. Using simulated data, we compare the imputation accuracy, power, and type I error rate of our proposed LA-aware approach to LA-unaware methods for performing TWASs. We show that CADET performs optimally in nearly all settings regardless of whether the genetic architecture of gene expression is dependent or independent of ancestry. We further illustrate CADET by performing a TWAS of 29 common blood biochemistry phenotypes within an admixed cohort from the UK Biobank and identify 18 hits unique to our LA-aware strategy with the majority of hits supported by existing GWAS findings.

## 1 INTRODUCTION

While genome-wide association studies (GWASs) have been largely successful in identifying genetic variants associated with a wide range of complex traits and diseases, many of the top variants lie in non-protein coding regions of the genome. It has been estimated that up to 90% of GWAS-identified single nucleotide polymorphisms (SNPs) are non-coding variants, and thus the biological mechanisms by which these variants exert their effects on a phenotype remain unclear^1^. Transcriptome-wide association studies (TWASs) seek to elucidate if the underlying biological mechanisms of such GWAS hits are due to complex gene regulatory effects. TWASs have achieved notable successes in the studies of various complex traits and diseases, including cancers, nervous and immune system disorders, anthropomorphic measurements, and many others^2–11^.

A typical TWAS assumes two datasets: a training dataset (usually of modest sample size) that possesses genotype and gene expression data from a tissue related to the outcome of interest, and a testing (GWAS) dataset that possesses genotype and outcome data but lacks expression data. TWASs involve a two-stage process. Stage I constructs a model on the training dataset to estimate the effects of genetic variants on expression levels of a target gene (expression quantitative trait locus weight, or eQTL weight). While traditional TWAS methods assume individual-level training data^7,10,12–15^, we note recent methods such as OTTERS allow more flexible use of summary-level eQTL training data for model fitting based on polygenic risk score (PRS) methodology^16^. After one fits training models for Stage I (using individual- or summary-level eQTL data), Stage II uses the resulting eQTL weights to impute genetically regulated gene expression (GReX) within subjects from the testing (GWAS) dataset and then tests for association between the imputed expression and the outcome of interest.

Existing TWASs generally assume that the training and test datasets are comprised from the same ancestrally homogenous population. Given the increased collection and analysis of multi-ancestry subjects for the testing GWAS data, there is need to expand the TWAS framework to handle genetic and genomic data from diverse groups. However, in order to develop methodology for multi-ancestry groups, we must consider that the underlying genetic architecture of gene expression may not be the same across populations and may differ by ancestral group^17^. Applied work has shown that gene expression prediction models trained in one ancestral population do not generalize well to other populations^18^. The gene expression prediction models used in such work are similar to PRS methods, which are known to have poor transferability across ancestral groups^19–23^. It is hypothesized that this poor portability may be a function of population-specific effect sizes, which can arise from a multitude of factors including differences in allele frequencies or linkage disequilibrium (LD) patterns across populations, as well as gene-gene or gene-environment interaction effects^24,26,28,30^.

The development of sufficiently powered TWAS methods for diverse ancestries is particularly important for admixed individuals, whose genomes are a unique mosaic of multiple continental ancestral groups. Admixed groups account for a large and growing proportion of the United States population, with more than 33.8 million people identifying as multiracial in the 2020 Census^25^. It has been common practice over the past few decades in GWASs to exclude admixed individuals from consideration due to their complex ancestral makeup, and as such, they are a historically underrepresented group in genetic studies. In response to this realization, researchers have recently considered the utility of including local ancestry information in variant-level genetic association analyses of admixed populations^27,29,31–39^. Building on these advancements, there further has been increased development of novel PRS approaches specifically designed for admixed populations that we can leverage for related TWASs^40–42^. For example, Marnetto et al. developed an ancestry-aware approach for PRS that first deconvoluted admixed haplotypes for a test subject and then computed the subject’s ancestry-specific components of the PRS using GWAS summary data from the appropriate reference ancestral populations. Authors demonstrated that this approach can not only yield improved phenotype predictability over standard methods but also an unbiased distribution of PRS in recently admixed populations^40^.

Given the existing evidence in the literature for differential eQTL architecture between admixed groups and more ancestrally homogenous subjects^17,43^, we propose CADET (<u>C</u>ombining <u>A</u>ncestry-<u>D</u>econvoluted <u>E</u>xpression in <u>T</u>WAS) for enhanced TWAS in admixed samples that leverages local ancestry information as well as summary-level eQTL data from multiple reference datasets of differing ancestry. In this method, we first apply the gene-expression training models used by OTTERS separately to each reference dataset. We then apply a local-ancestry deconvolution method to admixed test GWAS samples and, for a given gene, impute ancestry-specific partial GReX following the method of Marnetto et al. We combine the vectors of ancestry-specific GReX into an aggregate ancestry-aware measure of GReX. We can then test whether the aggregate ancestry GReX (as well as ancestry-specific GReX) are associated with outcome. We further combine the aggregate and ancestry-specific results together into an omnibus test using the Aggregated Cauchy Association Test (ACAT)^44^. We evaluate expression imputation accuracy and power of CADET in simulations. The procedure is well-calibrated under the null hypothesis of GReX-phenotype independence and achieves optimal power in the majority of genetic architecture settings. We conclude with an application of our method to 29 blood biochemistry phenotypes in admixed individuals of African and European ancestry in the UK Biobank and identify 18 significant gene-trait associations unique to our ancestry-aware strategy.

## 2 MATERIALS AND METHODS

### 2.1 Overview

The overarching goal of CADET is to impute gene expression in GWAS data of admixed individuals using summary-level eQTL data from individuals of the corresponding founder (continental) ancestral groups. Like most established TWASs, this involves two stages. Stage I focuses on using OTTERS to train models of predicted gene expression in each of the reference summary-eQTL datasets under consideration. Stage II then uses estimates from these trained models to construct both local-ancestry (LA) aware and LA unaware estimates of GReX (see overview of imputation workflow in Figure 1). We then test each LA aware and LA unaware GReX with an outcome of interest and aggregate results across tests into an optimal omnibus statistic based on the Cauchy distribution (see overview of downstream workflow in Figure 2). We describe how we construct our LA aware and unaware measures of GReX for admixed test subjects below.

**Figure 1.**
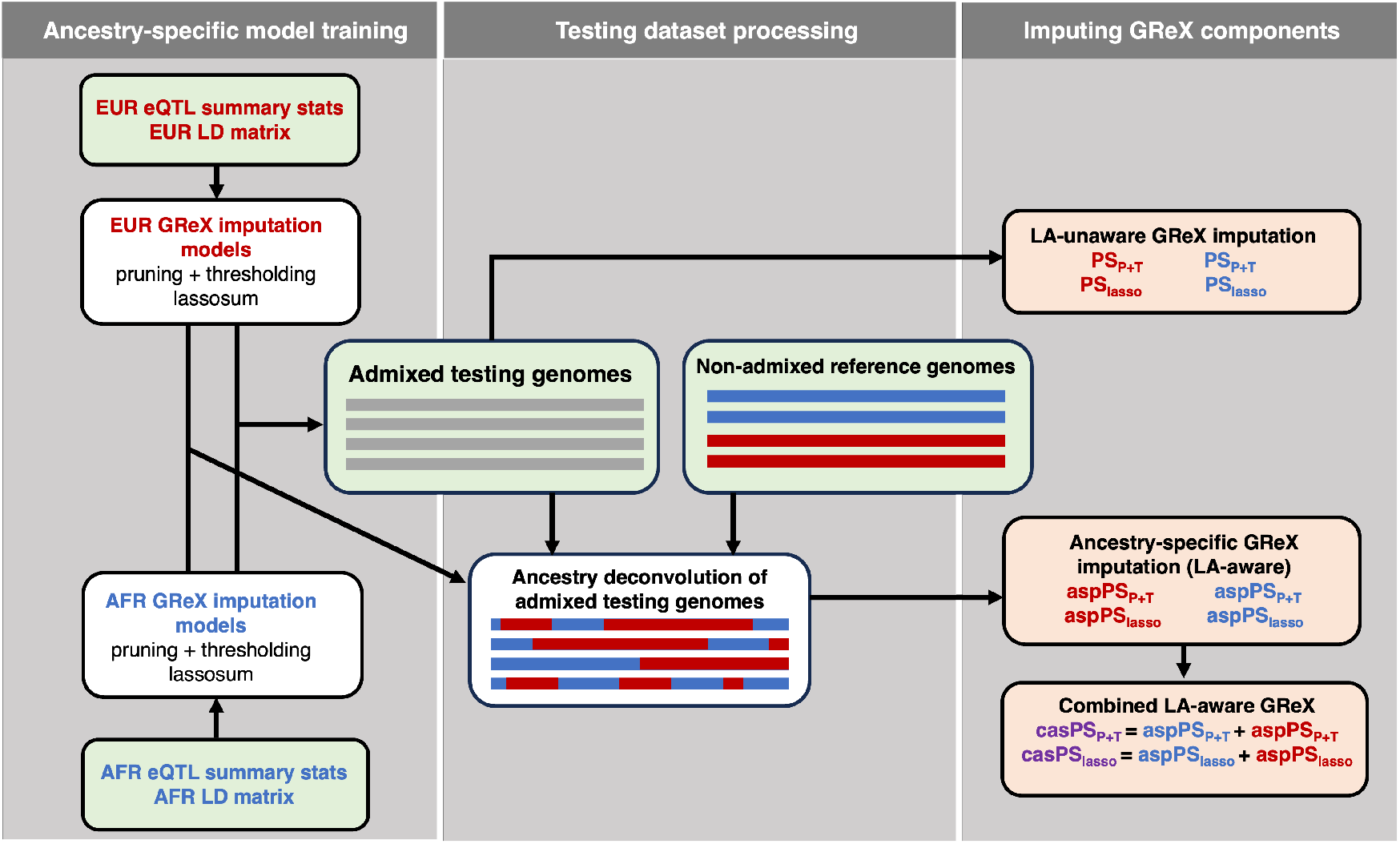
Overview of gene expression imputation in CADET. First, we use multiple external eQTL summary datasets from different ancestral groups and multiple polygenic risk score models (e.g., pruning and thresholding, lassosum) to train ancestry-specific GReX imputations models. Next, we use local ancestry (LA) deconvolution to parse the admixed genomes of our testing dataset (gray) into two groups defined by local ancestry (red and blue segments). For each gene, we define ancestry-specific genome subsets around the target gene based on LA assignment of each variant in the region. We then use our AFR- and EUR-specific GReX imputation models to compute several (LA-aware) GReX vectors using our deconvoluted admixed genomes, including the ancestry-specific PS (aspPS) and the sum of these scores constituting our combined LA-aware PS (casPS). We also use these imputation models to compute standard (LA-unaware) GReX vectors using the original non-deconvoluted admixed genomes. Green boxes indicate user-supplied datasets.

**Figure 2.**
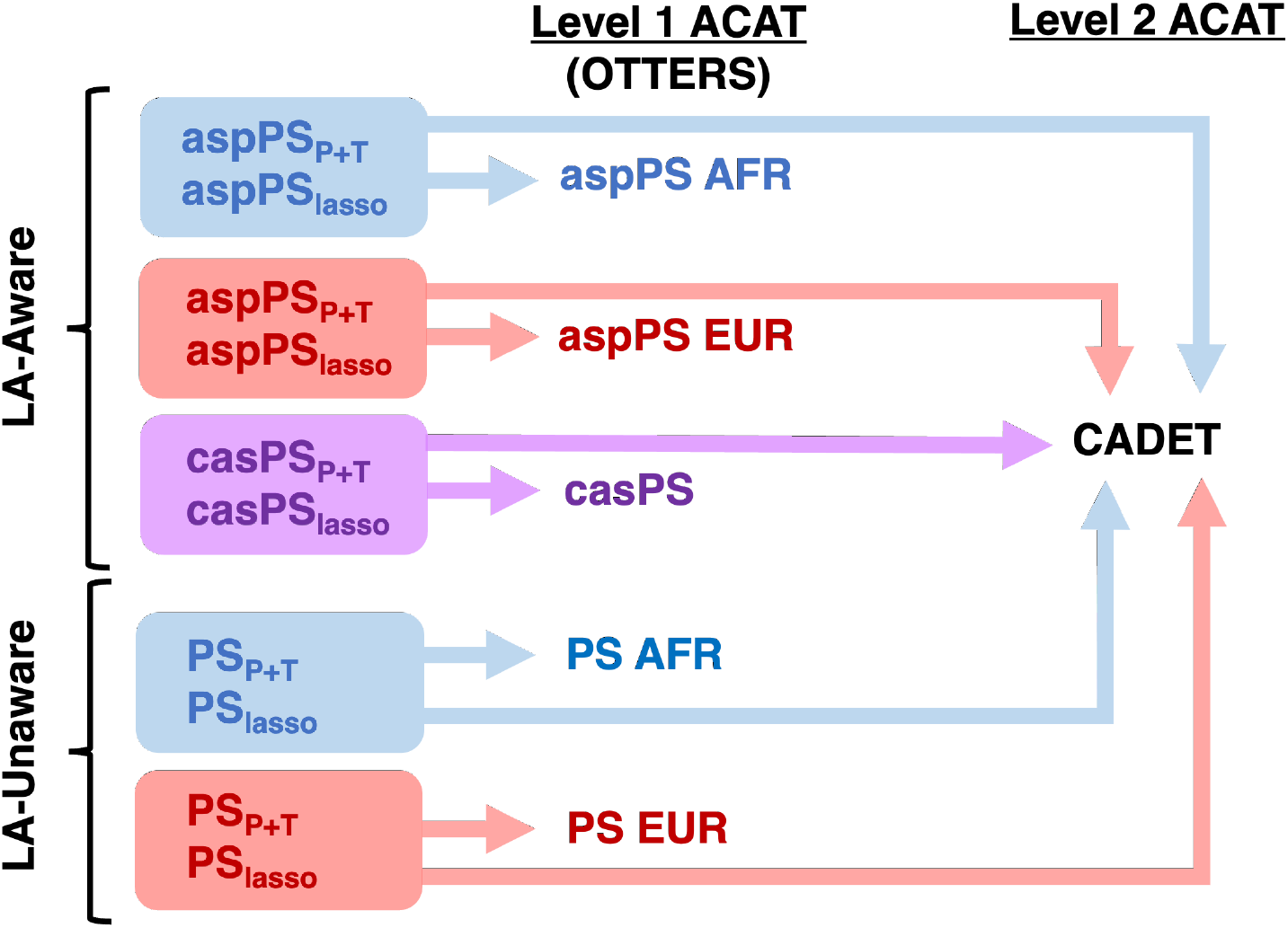
Overview of p-value aggregation in CADET. In this project, we compare the performance of CADET, which conducts Level 2 p-value aggregation across both PRS models (pruning/thresholding and lassosum), and imputation approaches (LA-aware and LA-unaware) to Level 1 p-value aggregation just across PRS models (the standard aggregation performed in OTTERS). We perform all p-value aggregation using the Aggregated Cauchy Association Test (ACAT).

### 2.2 Modeling Expression in Admixed Individuals

We first describe how we model gene expression in admixed subjects, which forms the backbone of our technique for estimating LA aware GReX in the test GWAS. Assume *N*_*adm*_ admixed individuals are two-way admixed, e.g., of African (AFR) and European (EUR) descent. For a given gene *g*, we assume there are *S* cis-eQTLs that are shared between the two ancestral groups and *U* cis-eQTLs that are unique to only one ancestral group. We define the total number of cis-eQTLs for a given gene as *V* = *S U*.

For 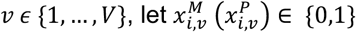 be the number of minor alleles for the *v*^th^ eQTL of subject *i* ( = 1, … , *N*_*adm*_) on the subject’s maternal (paternal) haplotype. Further, let 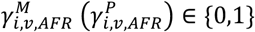 be an indicator variable that takes the value of 1 when the maternal (paternal) haplotype of subject *i* is of AFR origin and likewise define 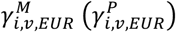 as a similar indicator variable for denoting when the maternal (paternal) haplotype of subject *i* is of EUR origin. Based on this notation, we can define 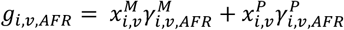 to represent the number of AFR-ancestry minor alleles of the *v*^th^ eQTL and likewise define 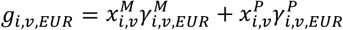 as the number of EUR-ancestry minor alleles at the eQTL for subject *i*. We can arrange these quantities across all subjects and all *V* eQTLs by creating matrices ***G***_***AFR***_ and ***G***_***EUR***_, each of dimension *N*_*adm*_ × *V*. In these matrices, we further assume each column has been centered to mean zero. We can therefore model the gene expression vector of our testing population as follows:

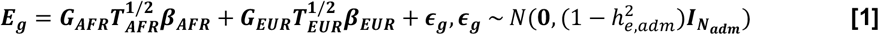

Here, ***E***_***g***_ is the *N*_*adm*_ × 1 vector of gene expression in our admixed testing population. ***T***_***AFR***_ and ***T***_***EUR***_ are the *V* × *V* diagonal scaling matrices with diagonal elements 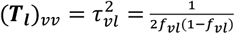, where _*vl*_ is the MAF of the *v*^th^ eQTL in reference population 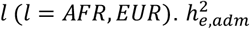 represents the gene expression heritability in our admixed subjects. ***β***_***AFR***_, ***β***_***EUR***_ ∈ ℝ^*V*^ are the vectors of causal eQTL effect sizes in each population. 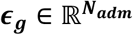 is the error vector, while 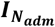 is the *N*_*adm*_ × *N*_*adm*_ identity matrix. Using these quantities, we can derive both the joint distribution of all eQTL effect sizes and an estimate for the heritability of gene expression in our admixed subjects (Supplemental Information).

### 2.3 Stage I: Reference Expression Model Training via OTTERS

Let us now assume, for our Stage I training datasets, we have summary-level eQTL data derived from *N*_*ref*_ individuals in each of the two reference populations that represent the source ancestries among our two-way admixed testing population. In other words, we require eQTL summary data from an AFR cohort and EUR cohort for imputing gene expression in African American individuals. We note that *N*_*ref*_ need not be the same for both AFR and EUR groups, but we assume so for ease of presentation.

In each homogeneous ancestral population (AFR, EUR), we can model GReX of each gene *g* as we did for our admixed subjects in 2.2. As above, expression is a function of the *V* cis-eQTLs with non-zero effect sizes in each population.

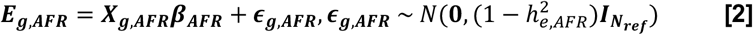

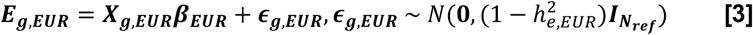

In the above equations, ***E***_***g***,***AFR***_ and ***E***_***g***,***EUR***_ are the *N*_*ref*_ × 1 vectors of gene expression in our AFR and EUR reference populations. ***X***_***g***,***AFR***_ and ***X***_***g***,***EUR***_ are the *N*_*ref*_ × *V* matrices of genotypes (0/1/2) with columns centered and standardized by minor allele frequency (MAF) in each population. ***β***_***AFR***_, ***β***_***EUR***_ ∈ ℝ^*V*^ are the vectors of ancestry-specific eQTL effect sizes. 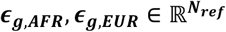 are the error vectors, while 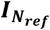 is the 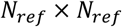 identity matrix. 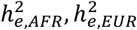 represent the gene expression heritability in each population.

A priori, we do not know the true *V* cis-eQTLs described in Equations 2 and 3. Instead, we attempt to model gene expression using eQTL summary information available from *J V* cis-SNPs found in and around gene *g*. That is, we have summary-level results (estimated effect sizes and p-values) from the following single-variant linear regression models for each cis-SNP *j* ∈ {1, … , *J*} within 1Mb of the transcription start site and end site of each gene *g*.

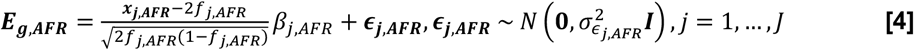

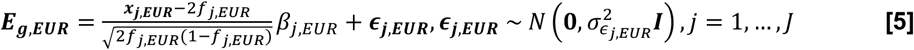

These linear regression models are fitted separately in each population *l*. In the above equations, ***x***_***j***,***AFR***_ and ***x***_***j***,***EUR***_ are the *N*_*ref*_ × 1 vectors of genotypes (0/1/2) for the *j*^th^ variant, *j* ∈ {1, … , *J*}, in the AFR and EUR reference populations, respectively. Similarly, we have error terms *ϵ*_***j***,***AFR***_ and *ϵ*_***j***,***EUR***_. *β*_*j*,*AFR*_ and *β*_*j*,*EUR*_ are the standardized marginal eQTL effect sizes for this variant. Derived from these fitted models, for each gene, we have summary statistics in the form of 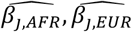 (the marginal least squares effect estimates) and corresponding p-values. These summarize the marginal association of variant *j* with the expression of the gene of interest in AFR and EUR populations. Using marginal estimated eQTL effect size vectors 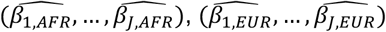 and corresponding marginal p-value vectors from the single-variant eQTL data, we use OTTERS to train PRS models to impute gene expression separately in each population. While the OTTERS pipeline includes multiple frequentist and Bayesian PRS methods as options, here we focus on two as illustrative examples. We briefly summarize the methodology of these two PRS models below:

#### Pruning and Thresholding (P+T)

This approach includes two steps: pruning, or clumping, of variants to exclude correlated SNPs, and thresholding to keep only those SNPs significantly associated with gene expression^45^. First, variants are filtered to include only those with p-values below a pre-specified threshold *P*_*T*_. Next, among these remaining variants, a set of pairwise-independent variants are selected such that linkage disequilibrium (LD) *R*^2^ among such variants is less than some pre-specified threshold *R*_*T*_, preferentially keeping those with the smallest p-value. These pruning and thresholding steps are performed in the OTTERS pipeline using PLINK 1.9^46^. For our analysis, we used thresholds *P*_*T*_ = (0.05, 0.001) and *R*_*T*_ = 0.99. We chose a liberal LD threshold as LD patterns differ considerably between populations, and OTTERS previously indicated that LD pruning did not significantly impact the performance of their method. We used the marginal standardized eQTL effect sizes from SNPs meeting these criteria to predict expression.

#### Lassosum

This method represents a summary-statistics-based version of the least absolute shrinkage and selection operation (lasso) pipeline; a penalized variable selection approach for dimension reduction with a large number of predictors (variants)^47^. Full details on lassosum are provided elsewhere^48^. In contrast to P+T methods, lassosum requires LD blocks from an external reference panel. LD blocks were pre-calculated using lddetect^49^ and data from 1000 Genomes (AFR and EUR populations, respectively)^50^. Similar to above, we performed LD clumping with *R*_*J*_ = 0.99 prior to training PRS via lassosum.

Using the two pruning and threshold approaches (P+T0.05, P+T0.001) and the lassosum approach, we have three sets of estimated eQTL effect size vectors for each training population for a given gene *g*: 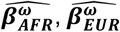 where ω ∈ {P+T0.05, P+T0.001, lassosum}.

### 2.3 Stage II: Imputing Expression in Admixed Individuals

#### LA Aware Approaches (aspPS, casPS)

As the true causal eQTLs in each ancestry are unknown, we propose to impute ancestry-specific components of GReX in the admixed testing set using the trained PRS models of gene expression in the two reference eQTL datasets. Let us assume that we have phased genotype information and have performed ancestry deconvolution of our admixed testing dataset haplotypes using existing methods like FLARE, RFMix, ELAI, or MOSAIC, among others^51–54^. For each PRS model considered ω ∈ {P+T0.05, P+T0.001, lassosum}, we use the estimated eQTL effect size vectors from Stage I, 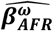 and 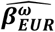, to impute **a**ncestry-**s**pecific **p**artial gene expression **P**R**S** (**aspPS**) as outlined in Figure 1 in the manner of Marnetto et al.^40^ We can then add together the AFR- and EUR-specific partial components of GReX to create a **c**ombined **a**ncestry-**s**pecific **P**R**S** (**casPS**).

Assume there are *J* estimated cis variants in the region of gene *g* and *N*_*adm*_ admixed testing subjects. Motivated by our underlying expression model for admixed subjects described in section 2.2, let **Γ**^***M***^ (**Γ**^***P***^) be a *N*_*adm*_ × *J* matrix where the *(i*,*j)* element equals 1 if individual *i* has an AFR allele at *j*^th^ SNP on his/her maternal (paternal) haplotype and 0 otherwise. We further define **X**^***M***^ (**X**^***P***^) as the *N*_*adm*_ × *J* ancestry-standardized maternal (paternal) haplotype (0/1) matrix with 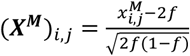 . Here, 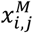 is the minor allele count of the *i*^th^ individual at the *j*^th^ variant on the maternal haplotype and *f* is the MAF of that variant in EUR or AFR, depending on the local ancestry of that SNP. Let 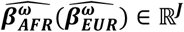 be the estimated eQTL effect size vector from a given PRS method ω using reference AFR (EUR) eQTL data. In CADET, we impute the AFR component of GReX 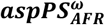 and the EUR component of GReX 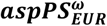 as follows:

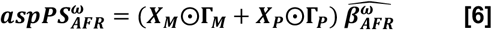

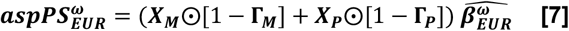

In the above equations, the symbol ⨀ indicates element-wise multiplication of matrices. For each PRS model ω, we then impute the total GReX in our admixed testing samples as the sum of these two ancestry-specific components:

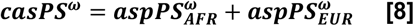

#### LA Unaware Approaches (PS)

We can also compare the performance of the ancestry-aware approaches to imputing GReX (***aspPS***_***AFR***_, ***aspPS***_***EUR***_, ***casPS***) to LA unaware approaches that simply apply the standard PRS methodologies to admixed individuals. For this, let ***X*** be the *N*_*adm*_ × *J* standardized (columns centered and scaled to unit variance, not ancestry-specific standardization) matrix of minor allele counts (0/1/2). We can impute GReX in our admixed testing data as:

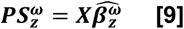

Here, 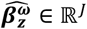 is the estimated eQTL effect sizes vector from a given PRS method ω for a given set of eQTL summary statistics *z*. This standard approach could use the same reference population eQTL summary data as those used in the LA-aware approaches above (*z* = AFR, EUR), or it could be trained using summary eQTL data from a sample of admixed individuals that is independent of our testing dataset (*z* = ADMIX).

## 2.5 Stage II: Gene-Trait Association Test

We assume that our phenotype vector 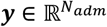, for whom we want to estimate the association with GReX, has already been adjusted for the effects of important non-genetic confounders. Once we have our all of our vectors of imputed gene expression in our admixed testing dataset 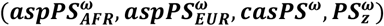 for *ω* ∈ {P+T0.05, P+T0.001, lassosum} and *z* ∈ {AFR, EUR, ADMIX}, we perform simple linear or logistic regression of ***y*** on each of the aspPS, casPS, and PS imputed GReX vectors individually. For each ω and *z*, we thus obtain an ordinary least squares p-value. Using the approach employed by OTTERS^16^, CADET then combines these p-values in multiple ways (as outlined in Figure 2) using the Aggregated Cauchy Association Test (ACAT)^44^. First, we can aggregate across ω (our PRS models) to generate one p-value for each of our LA-aware approaches (*aspPS*_*AFR*_, *aspPS*_*EUR*_, *casPS*), as well as our LA-unaware approaches (PS). We refer to this as Level 1 aggregation, and this is the typical p-value aggregation performed in OTTERS. For example, to obtain the Level 1 p-value for casPS, let *p*_*ω*_ be the p-value from the simple regression of ***y*** on ***casPS***^*ω*^ for ω ∈ {P+T0.05, P+T0.001, lassosum}. We then calculate the Level 1 p-value for casPS (*p*_*casps*_) as

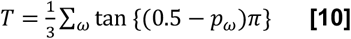

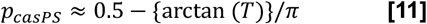

Here, *T* is the ACAT test statistic, which is assumed to follows a Cauchy distribution. In (10) and (11), we assume the weights (1/3) to be equal across all PRS methods. We can then repeat this Level 1 aggregation process to further calculate 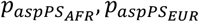, and 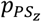.

As shown in Figure 2, CADET then performs a second round of p-value aggregation (which we refer to as Level 2 aggregation) that combines all LA-aware p-values 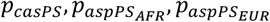 from Level 1 analyses and can further incorporate LA-unaware p-values 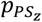, if desired. We combine such p-values using the same ACAT framework outlined for Level 1 analysis as shown in equations (10) and (11).

## 2.6 Simulations

To evaluate the accuracy of our proposed ancestry-aware gene expression imputation approach, we performed extensive simulations. First, we chose our simulated admixed testing dataset to be two-way African and European recently admixed individuals. We used the 1000 Genomes (1KG) Phase 3 biallelic data in GRCh38 as source genomes to simulate multiple sets of admixed genomes^50^. We specifically chose CEU (Utah Residents with Northern and Western European Ancestry) as the European source population and YRI (Yoruba in Ibadan, Nigeria) as the African source population. To ensure our source populations were large enough for sampling haplotypes to generate our admixed testing set, we first expanded the YRI and CEU populations to size 10,000 each using admix-kit^55^. This tool generates additional sets of 1KG populations using the HAPGEN2 framework^56^.

We selected one gene, *ABCA7*, on chromosome 19 (GRCh38 1040101:1065571) as our target gene for simulations, as this gene has been used previously for the simulation stage of TWASs^13^. We assumed a window of 1Mb upstream and downstream as the simulation region (66,358 variants). We generated different training datasets in which we first simulate gene expression and then subsequently calculate eQTL summary data. We generated AFR and EUR training datasets by randomly selecting 500 of our expanded sample of YRI and CEU subjects, respectively. We further generated several admixed training datasets of size 500 using the expanded sets of 10,000 YRI and CEU source haplotypes and the tool haptools^57^. We simulated three admixed training datasets according to different admixture generation parameters. First, we assumed an initial realistic African American demographic model of one pulse of admixture 10 generations ago with 80% contribution from YRI and 20% from CEU (ADMIX 80 10g). We also considered two additional admixed training sets; the first assumed 5 admixture generations plus an initial AFR population frequency of 80% (ADMIX 80 5g) while the other assumed 5 admixture generations plus an initial AFR population frequency of 50% (ADMIX 50 5g). We considered these additional training sets as previous work in this field has shown that power of TWAS association tests can be negatively impacted by not only differing training and testing populations (e.g., EUR vs. admixed), but it can also vary according to admixture generation parameters in the admixed testing genomes, such as the initial AFR population frequency^18^.

To generate our admixed testing datasets, we again assumed the realistic African American demographic model of 10 admixture generations with 80% contribution from YRI (AFR) and 20% from CEU (EUR). These admixture generation settings for our testing dataset match those used to generate the ADMIX 80 10g training dataset. These simulation settings are similar to those used in previous methodological work in admixed individuals^27^. We simulated a total of 10,000 individuals to serve as our pool of admixed testing samples.

Using the simulated haplotypes and genotypes from the non-admixed reference AFR/EUR populations, we next simulated gene expression. In our training datasets (admixed and reference AFR/EUR of size 500), we first simulated gene expression according to the models described in Sections 2.2 and 2.3. We varied gene expression heritability in the AFR population 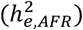 and in the EUR population 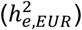 among {0.1,0.2}, excluding the limited utility scenario where both are 0.1. We also varied the number of eQTLs (*V*) in both populations among {2,10,100}, the proportion of eQTLs that overlap between AFR and EUR populations (OP *= S/V*) among {0.5, 1}, and the correlation of effect sizes among shared AFR and EUR eQTLs (*ρ*) among {0.5, 1}. We also ensured that all SNPs selected as eQTLs in each population have MAF > 0.05 in the corresponding training datasets. In total, we have 36 combinations of expression simulation parameters.

Since all settings consider overlap in eQTLs across ancestries, we took care when simulating correlated eQTL effect size vectors in each ancestry. Using the notation of Equations 2 and 3, we first randomly drew a temporary 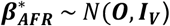 and calculated the scale factor 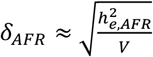. To simulate eQTL effect sizes in EUR, note that the first *S* are correlated with the first *S* elements of the effect size vector in AFR. Thus, we sampled a temporary 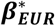 as below:

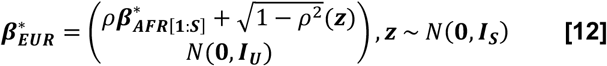

Let the scale factor for EUR be 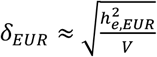 . Thus, for each simulation, the eQTL effect size vectors achieving the desired 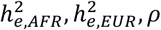 and OP are simply 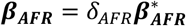 and 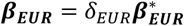. We then used these ancestry-specific eQTL effect size vectors to simulate gene expression in our reference AFR/EUR training sets and, also, our admixed training/testing sets according to Equation 1.

Next, for each simulated training dataset (reference AFR/EUR and our independent admixed training samples), we calculated single-variant eQTL summary statistics and used the OTTERS pipeline to train PRS models. For each dataset, we retained summary data for only those variants with MAF > 0.05 in the corresponding sample. Using these eQTL summary statistics, we then imputed gene expression in our admixed testing set in the manner described in Section 2.3. We compared the imputation R^2^ (squared correlation between imputed and true expression in our admixed testing set) between our proposed LA-aware approaches 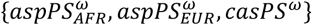, ω ∈ {P+T_0.05, P+T_0.001, lassosum}, and LA-unaware approaches *PS*_*ω*_ for the AFR, EUR, ADMIX 80 10g, ADMIX 80 5g, and ADMIX 50 5g training datasets. For each round of GReX imputation using LA-aware approaches, we also performed another round of imputation where we assumed that 10% of the cis-variants in the region on both haplotypes of each individual in the testing set had incorrect local ancestry information.

Next, we performed power and type I error simulations for the Stage II test of GReX-trait association. We simulated the trait according to following equation where we assume the trait is a function of the total gene expression and is not a function of the ancestry-specific components of gene expression in Equation 1.

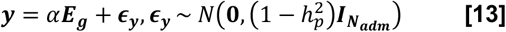

For both our power and type I error simulations, we calculated the Level 1 ACAT p-values 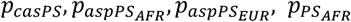 and 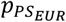. We then also calculated CADET’s Level 2 ACAT p-values that aggregated over various combinations of these Level 1 ACAT p-values. For our power simulations, we considered such that 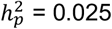. For our null simulations to evaluate type I error rate, we assume this value was 0. For each of the 1,000 simulations performed for each of the 36 combinations of expression simulation parameters, we performed 10 trait simulations per 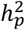 Thus, for each expression simulation parameter combination, we performed 10,000 power and 10,000 type I error simulations.

## 2.7 CADET Analysis of Admixed Subjects from UK Biobank

### UK Biobank Data

To assess the utility of CADET for TWASs in practice, we obtained individual-level genotype and phenotype data from admixed individuals in the UK Biobank (UKB), which is a large-scale biomedical database housing data collected from approximately 500,000 individuals across the UK. To best mimic the settings of our simulation study, we elected to focus our analysis on two-way admixed individuals of African and European ancestry in the UKB. In order to identify these individuals comprising our testing dataset, we first performed preliminary subject filtering. Specifically, we excluded subjects who had subsequently withdrawn from the study, those who were marked as “redacted”, and those who were marked as outliers based on pre-calculated metrics of heterozygosity and missing rates. We also excluded subjects with putative sex aneuploidy, those with high pre-calculated estimates of relatedness, and those whose genetically-inferred gender differed from their submitted gender. We then removed individuals falling in the “White British subset.” These individuals were previously identified by a combination of both self-reported ancestry and genetic PCs. We also excluded individuals who have missing self-reported ethnicity or whose self-reported ethnicity fell among the following: White, Irish, British, Any other white background. Following this, 27,491 subjects remained.

We then performed principal component analysis (PCA), projecting these filtered UKB individuals onto genetic PCs anchored in 1000 Genomes (1KG) data^50^. For this, we first subset 1KG genotype data to include unrelated individuals from the following populations: African (ACB [African Caribbeans in Barbados] and ASW [Americans of African Ancestry in SW USA] excluded) (503), Admixed American (347), East Asian (504), European (503), and South Asian (489). We restricted variants to non-ambiguous SNPs, those found in the UKB GWAS data, those with MAF > 0.05 in each population, and those with Hardy-Weinberg equilibrium (HWE) p > 1E-6. We then pruned remaining variants (window size 1000 bp, step size 50 variants, R^2^ threshold 0.1). Using the loadings for the top 10 PCs trained in 1KG samples, we projected the UKB self-reported non-White individuals (27,491) into this space (Figure S1). Following the approach of Atkinson et al.^27^, using the 1KG data, we then trained a random forest classifier to predict continental ancestry (1KG population) from the top 10 PCs. We then applied this random forest model to our UKB sample. We excluded any individuals with <50% estimated probability of African ancestry. Using the top 3 PCs in the 9,187 meeting these criteria, we constructed a 95% ellipsoid along the African-European cline (Figure S2). We kept the 8,752 UKB individuals lying within the ellipsoid. Finally, we excluded two additional subjects with self-reported Asian/Asian British and White and Asian ethnicity. The remaining 8,750 individuals made up our final two-way AFR and EUR admixed testing dataset.

Genotype data on these subjects was generated using either the UK BiLEVE or UK Biobank Axiom arrays. Prior to imputation, we followed the pre-imputation quality control pipeline provided at https://www.well.ox.ac.uk/~wrayner/tools/\#Checking, using variant data from TOPMed Freeze3a on GRCh37/hg19. We performed imputation, liftover to GRCh38, and phasing using the TOPMed Imputation Server^58–60^. Next, we prepared our AFR and EUR reference data for local ancestry deconvolution of our UKB genotypes. First, we imputed missing genotypes in the 1KG Phase 3 biallelic phased GRCh38 data of AFR and EUR subjects using BEAGLE, again excluding ASW and ACB^61^. Using this as our reference population genotype data, we performed local ancestry inference in our admixed UKB testing set using FLARE^51^. We assumed 10 generations since admixture (10 admg).

### eQTL Reference Summary Data

For our European (EUR) reference eQTL dataset, we downloaded cis-eQTL summary statistics in whole blood from the GTEx V8 dataset (dbGaP phs000424.v8.p2), where cis-eQTL analysis was performed in 570 European-American subjects. To briefly summarize the analysis performed, authors adjusted RNA sequencing for the effects of top 5 PCs, top 60 PEER factors, sequencing platform (Illumina HiSeq 2000 or HiSeq X), sequencing protocol (PCR-based or PCR-free), and sex. For the eQTL analysis, they then restricted genes to those with >0.1 TPM and ≥ 6 reads in at least 20% of the data samples, and they normalized expression vectors using TMM^62^ and inverse normal transformation. SNPs from WGS data with MAF ≥ 1% were retained. Authors performed single-variant cis-eQTL analysis using FastQTL^63^ and a 1MB window from the transcription start site of each gene.

For our African (AFR) reference eQTL dataset, we used publicly-available whole-blood cis-eQTL summary statistics from a subset of high-African ancestry admixed individuals^17^. In this study, authors mapped ancestry-specific gene expression signatures in 2,733 individuals (African American, Puerto Rican, and Mexican American) from the Genes-Environments and Admixture in Latino Asthmatics (GALA II) study and the Study of African Americans, Asthma, Genes, and Environments (SAGE). Authors obtained paired RNA sequencing data and WGS data. RNA and WGS data processing are described elsewhere^17^. Authors used CEU and YRI HapMap reference genotypes, as well as Indigenous American ancestry reference genotypes, to estimate global measures of ancestry with ADMIXTURE^64^. Authors defined a high global African ancestry subset as those with >50% estimated global African ancestry (721). They then performed ancestry-specific eQTL analysis in these subjects using a 1Mb cis-window from the transcription start site of each gene and FastQTL^63^. Analyses adjusted for expression for age, sex, asthma status, top 5 PCs, and 60 PEER factors.

### TWASs of Blood Biomarkers in UKB

We used CADET to perform TWASs of 29 widely collected blood biomarkers in our UKB testing dataset. For each blood biomarker, we first log-normalized raw trait measurements and then we obtained covariate-adjusted phenotypes by taking the residuals of linear regression models of each log trait on the top 20 PCs, sex, age at recruitment, and smoking status (prefer not to answer, never, previous, current). Using our AFR and EUR reference eQTL summary data described above, we first removed ambiguous SNPs (A/T,T/A,G/C,C/G) and only kept eQTL data from SNPs with MAF > 0.05 in each respective sample. Next, we trained AFR and EUR ancestry-specific PRS models of gene expression using OTTERS and P+T0.001, P+T0.05, and lassosum models. We then imputed gene expression in UKB samples using LA-aware approaches (casPS, aspPS) and standard PS approaches (PS AFR, PS EUR). We concluded our analyses by performing simple linear regression analysis for the association of each imputed gene expression vector with each of the 29 adjusted blood biomarker traits. We considered multiple Level 2 p-value aggregation approaches via ACAT.

## 3 RESULTS

### 3.1 Expression Imputation Accuracy

We first compared GReX imputation accuracy of proposed LA aware approaches to LA unaware approaches assuming *N*_*adm*_ = 10000 simulated admixed individuals (10 admixture generations, 80% initial contribution of AFR [1KG YRI] haplotypes, 20% contribution of EUR [1KG CEU] haplotypes). We computed CADET’s LA-aware measures of GReX using simulated eQTL summary data from a reference AFR and a reference EUR sample (each *N*_*ref*_ = 500) and computed the LA-unaware GReX using each of these reference samples (AFR PS, EUR PS). We also computed additional LA-unaware GReX assuming eQTL summary datasets from independent simulated admixed samples of varying admixture parameters. We considered admixed reference samples with the same configuration as our test GWAS (10 admg, 80% initial AFR contribution) as well as other admixed reference samples that differed from the test GWAS either in admg (5 versus 10), AFR contribution (50% versus 80%), or both to examine the impact of mismatch between admixed training and test datasets on imputation accuracy.

Assuming an admixed testing sample size of 10,000, we present the squared correlation (R^2^) between imputed GReX and true gene expression assuming gene expression heritability of 0.2 in both Africans and Europeans 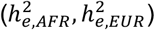 under the existence of two true eQTLs (Figure 3) or 10 true eQTLs (Figure S3). We provide imputation R^2^ results for other expression heritability and eQTL architecture settings in Figures S3-S5. From these figures, we see a few general trends. Across all LA-aware and LA-unaware imputation approaches, as well as across all PRS methods (P+T0.001, P+T0.05, lassosum), we see higher R^2^ values for the sparse eQTL scenario (2) compared to the scenarios where the number of causal eQTLs is 10 or 100. There may exist other PRS approaches (PRScs, for example^65^) that may perform better for the scenario in which we expect larger number of true eQTLs.

**Figure 3.**
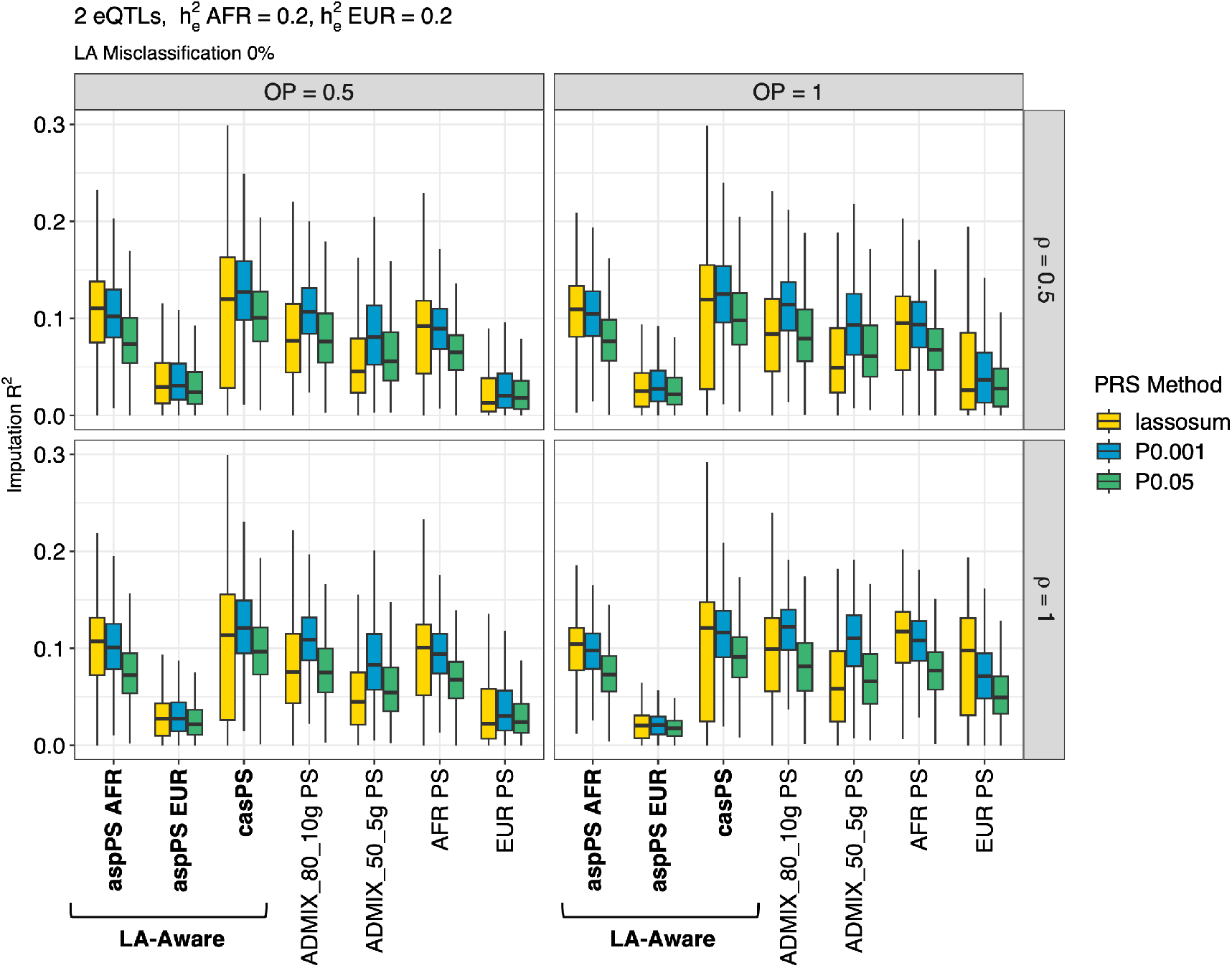
Gene expression imputation accuracy in 10,000 admixed testing samples (10 admixture generations, 80% initial contribution from AFR, 20% initial contribution from EUR) for expression heritability 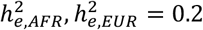 and 2 eQTLs. Vertical panels indicate the proportion of eQTLs that overlap (OP) between AFR and EUR ancestries, and horizontal panels indicate the correlation in eQTL effect sizes for shared eQTLs between the two ancestral groups (*ρ*). The x-axis shows the GReX imputation approach, including our proposed local-ancestry aware methods (aspPS, casPS) and standard PRS imputation approaches (PS). For ancestry-aware methods, we assume no local ancestry misclassification. Whiskers of boxplot extend to maximum/minimum point that is less than 1.5*IQR from the third/first quartiles.

Figures 3 and S3 also demonstrate that the accuracy of our proposed casPS approach tends to be slightly higher than the optimal standard PS approach using admixed eQTL summary data from a perfect ancestry-matched admixture cohort (ADMIX 80 10g PS) when the eQTLs are not identical between AFR and EUR or when the correlation of shared eQTL effect sizes is less than 1. In other words, these simulations suggest that even if we had access to eQTL data from a cohort exactly matched for ancestry to our admixed testing cohort, we would still achieve as high or higher imputation accuracy using casPS with reference population eQTL datasets. In the scenario where eQTL architecture for the given gene is identical between AFR and EUR populations (OP = 1, *ρ* = 1), the imputation R^2^ appears quite similar between the casPS LA-aware approach and ADMIX 80 10g PS. Furthermore, across all settings, we observe that R^2^ decreases with increasing differences in admixture generation parameters between training and testing datasets. As the number of admixture generations decreases (10 to 5) and the proportion of initial AFR donors decreases (80% to 50%) within the training datasets, we generally see less successful GReX imputation in the test dataset generated assuming 10 admixture generations and initial AFR population frequency of 80%.

These initial results assume correct inference of local ancestry in the admixed test samples. To assess the impact of LA misclassification on imputation accuracy, we conducted additional simulations where 10% of the eligible cis-SNPs of the test gene with MAF > 0.05 in the corresponding training datasets (Ref AFR or Ref EUR) had incorrect LA tags (AFR or EUR). We further assumed misclassification of these SNPs on both maternal and paternal haplotypes. We note this assumption is likely on the high end for misclassification rates, as popular local ancestry imputation algorithms (e.g., FLARE^51^, MOSAIC^54^, RFMix^52^) have high imputation accuracy for reasonably sized training panels used to infer LA^51,66^. Nevertheless, as shown in Figures S6-S8, we do not see a marked drop in the quantiles of imputation R^2^ of our proposed LA-aware GReX imputation approaches (aspPS AFR, aspPS EUR, casPS) across PRS models.

### 3.2 Type I Error Rate

Next, we assessed the type I error rate of the Level 1 and Level 2 TWAS association tests. For these simulations, we assumed a null association between true gene expression and each simulated phenotype, i.e., a phenotypic heritability 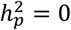. As the type I error rates did not differ dramatically by the gene expression simulation parameters (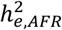 , 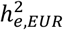 , *ρ*, OP, number of eQTLs), we present quantile-quantile (QQ) plots for p-values across each of the 36 simulation settings, corresponding to a total of 36,000 null simulations per plot.

As we see, the p-values resulting from Level 1 aggregation across PRS models (P+T0.001, P+T0.05, lassosum) for the LA-aware approaches (aspPS AFR, aspPS EUR, casPS) show the expected distribution under the null when we assume no LA misclassification (Figure S9) and 10% LA misclassification (Figure S10). Similarly, the Level 1 p-value aggregation for the standard imputation approaches using reference eQTL data (PS AFR, PS EUR) also appear to maintain appropriate rates of type I error. To assess whether we see any inflation of our gene-trait association p-values at the second level of p-value aggregation, i.e., aggregating Level 1 p-values for our LA-aware approaches and/or standard PS approaches, we constructed a second set of QQ plots in Figure S11. Again, we see the expected distribution under the null even when we combine aspPS and casPS p-values with those from the reference-derived PSs (AFR PS, EUR PS) and, further, with the p-values from the independent admixed-derived PSs (middle and right panels, respectively). This applies to both assumptions of 0% (top row) or 10% LA misspecification (bottom row).

### 3.3 Power

Simulation results comparing the performance of our proposed LA-aware approaches to the competing standard GReX imputation approaches under the assumption of a true gene-trait association are summarized in Figure 4 (2 eQTLs) and Figure 5 (10 eQTLs). These figures reflect normally distributed traits, a testing set sample size of 10,000, significance level α= 2.5E-6, and no LA misspecification. When comparing the performance of the Level 1 approaches (aggregation across PRS), our proposed LA-aware casPS has higher or equal power compared to any of the LA-unaware standard PS approaches, including that trained using a perfectly matched admixed sample, when gene expression genetic architecture is different between populations. Of note, the power of Level 1 casPS is also higher than that of a LA-unaware method with a perfectly matched admixed sample (ADMIX 80 10g) when the gene expression genetic architecture is exactly the same between AFR and EUR groups (OP = 1, *ρ* = 1). Furthermore, across all simulation settings, results show that Level 2 aggregation of LA-aware and LA-unaware methods using CADET achieved greater power than any individual Level 1 LA-unaware method. In Figures S12 (2 eQTLs) and S13 (10 eQTLs), we illustrate the power of the LA-aware approaches assuming a high rate of LA misclassification (10%) for cis-SNPs in the gene region. With the exception of the scenario in which we have a higher level of gene expression heritability in Europeans 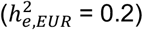 and lower level of expression heritability in Africans 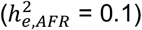 and assume 10 eQTLs, we still observe the Level 2 ACAT LA-aware approach of CADET achieving highest power. For the anomalous setting mentioned, we note that gene-expression imputation accuracy here is quite low across the board (Figure S5), and thus power is subsequently low for all LA-aware and LA-unaware approaches. This is similar to the scenario where we assume 100 eQTLs for the gene under study. We have comparatively less accurate GReX imputation, and therefore it is unsurprising that we see low power in the downstream association tests across all methods (Figures S14-S15).

**Figure 4.**
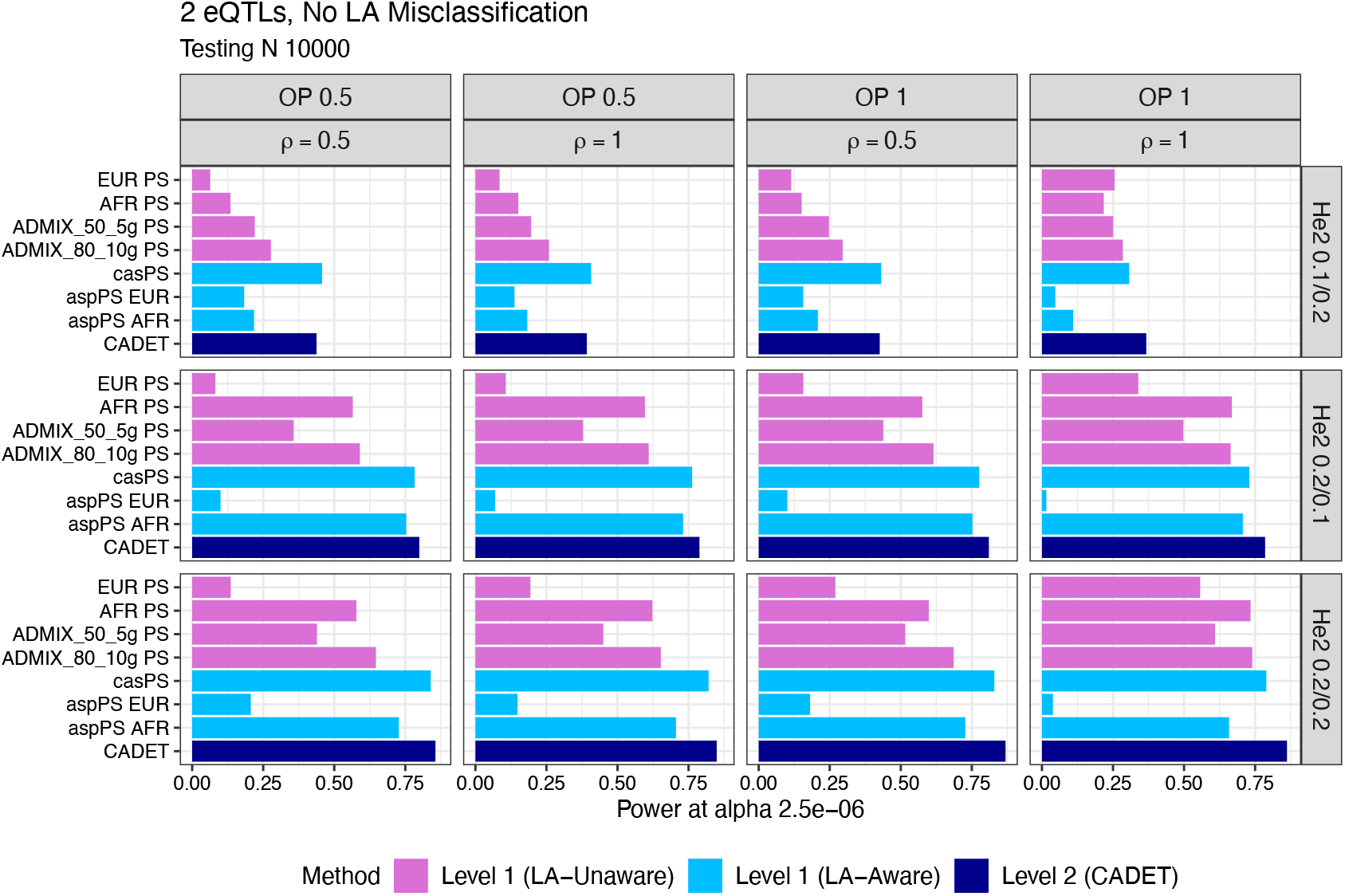
Power of gene-level association tests of imputed GReX vectors and simulated trait at significance level 2.5E-6. Here, we assume a phenotypic heritability of 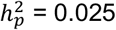, 2 eQTLs, no local ancestry (LA) misclassification for LA-aware approaches, and a testing dataset sample size of 10,000. Vertical panels indicate the overlapped proportion of eQTLs between AFR and EUR ancestries (OP) and the correlation of eQTL effect sizes for shared eQTLs (*ρ*). Horizontal panels indicate the gene expression heritability in AFR and EUR ancestries ( ^2^ AFR/EUR). Pink bars indicate the power of LA-unaware GReX imputation approaches, with p-values aggregated across the three PRS models. Light blue bars indicate LA-aware approaches with Level 1 p-value aggregation by ACAT. Dark blue bars indicate the power of Level 2 (CADET) analysis aggregating p-values from all LA aware and LA unaware approaches.

**Figure 5.**
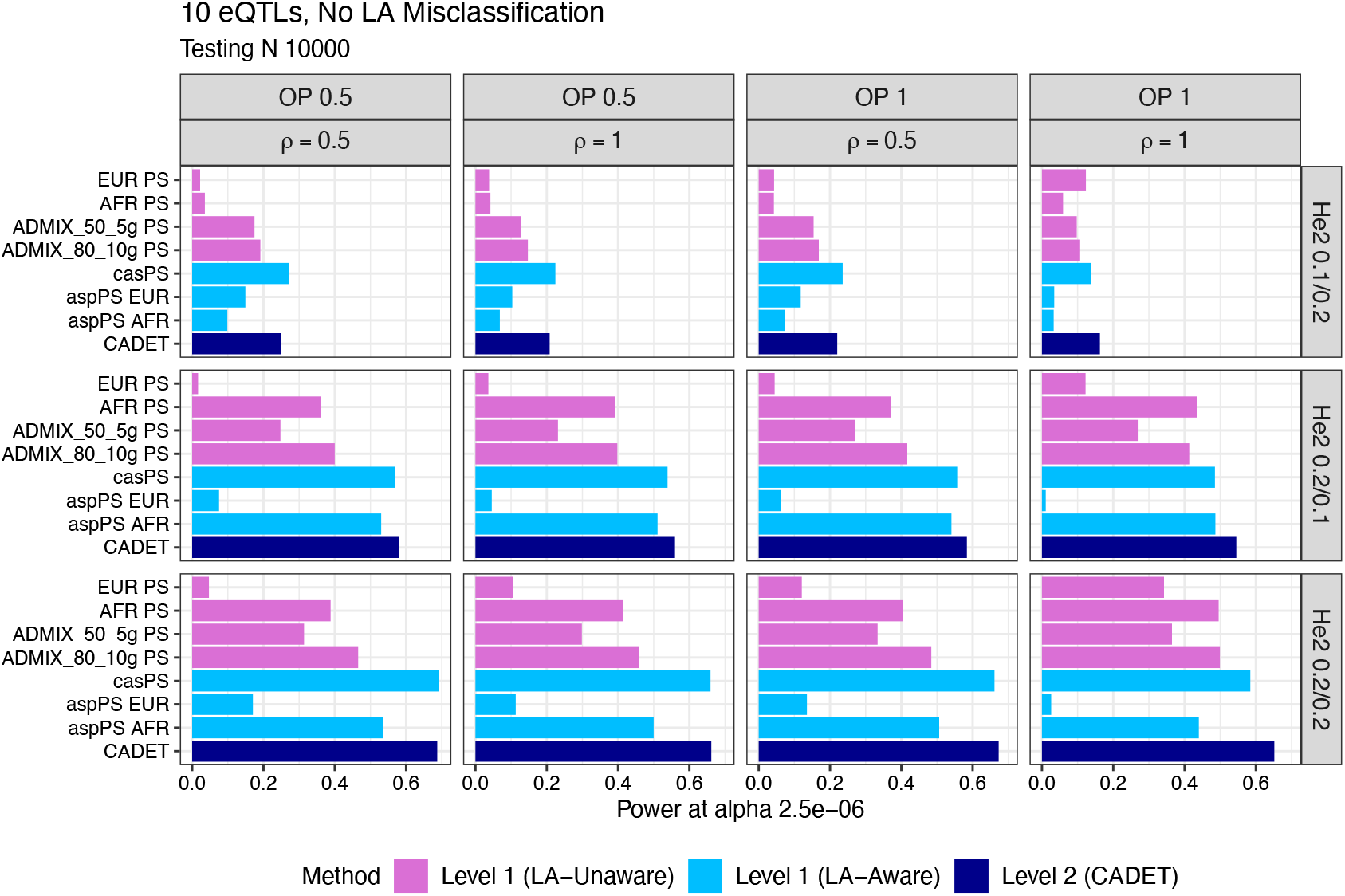
Power of gene-level association tests of imputed GReX vectors and simulated trait at significance level 2.5E-6. Here, we assume a phenotypic heritability of ^2^ = 0.025, 10 eQTLs, no local ancestry (LA) misclassification for LA-aware approaches, and a testing dataset sample size of 10,000. Vertical panels indicate the overlapped proportion of eQTLs between AFR and EUR ancestries (OP) and the correlation of eQTL effect sizes for shared eQTLs (*ρ*). Horizontal panels indicate the gene expression heritability in AFR and EUR ancestries (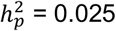 AFR/EUR). Pink bars indicate the power of LA-unaware GReX imputation approaches, with p-values aggregated across the three PRS models (ACAT Level 1). Light blue bars indicate LA-aware approaches with Level 1 p-value aggregation by ACAT. Dark blue bars indicate the power of LA-aware approaches, aggregating both PRS p-values and the resulting p-values of casPS, aspPSs (AFR and EUR), and standard PSs trained in the two AFR/EUR reference populations (ACAT Level 2).

### 3.4 CADET Analysis of UK Biobank

We applied CADET to detect genes associated with blood biomarkers by way of genetically-regulated transcriptional activity in a subset of 8,750 two-way African/European admixed subjects in the UK Biobank. We first performed LA deconvolution using FLARE and reference genotypes from AFR and EUR cohorts from 1000 Genomes. Through this, we estimated an overall global (genome-wide) proportion of AFR genotypes of 86.6% and an estimated proportion of EUR genotypes of 13.4%. Using our two sets of European-derived and high-African ancestry-derived eQTL summary data in whole blood, we trained our AFR and EUR GReX imputation models for 15,093 genes. Specifically, we used lassosum and pruning and thresholding models, keeping 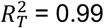 as in the simulations and filtering by p-value thresholds of *P*_*T*_ = 0.05, 0.001. For each PRS model, we then imputed GReX in our admixed UKB testing set in the standard OTTERS approach separately using our AFR-trained models (PS AFR) and EUR-trained models (PS EUR). We also imputed GReX in our proposed LA-aware approach using both sets of PRS models and our inferred LA information. We calculated two ancestry-specific components of gene expression (aspPS AFR, aspPS EUR) and one vector representing the sum of these two components (casPS).

We tested for the association of each of the vectors of imputed GReX with 29 blood biomarkers: 4 bone and joint traits (alkaline phosphatase, calcium, rheumatoid factor, vitamin D), 8 cardiovascular traits (apolipoprotein A and B, C-reactive protein, cholesterol, HDL cholesterol, LDL cholesterol, lipoprotein A, triglycerides), 2 diabetes-related traits (glucose, HbA1c), 3 hormone traits (insulin-like growth factor 1 [IGF-1], sex hormone binding globulin [SHBG], testosterone), 6 liver-related traits (alanine aminotransferase, albumin, aspartate aminotransferase, direct bilirubin, gamma glutamyltransferase, total bilirubin), and 6 traits related to renal function (creatinine, cystatin C, phosphate, total protein, urate, urea). For each approach (PS AFR, PS EUR, aspPS AFR, aspPS EUR, casPS), we calculated a Level 1 p-value aggregated across PRS models. We also aggregated across a second dimension beyond PRS model to calculate two Level 2 p-values (casPS+aspPS, casPS+aspPS+PS [CADET]).

We assessed significance at a Bonferroni-adjusted level of 0.05/(29*15093) = 1.14E-07, adjusting for the total number of phenotypes and number of genes considered. Across all tests and imputation approaches, we identified 384 significant gene-trait associations. While 384 significant associations were found, most gene-trait associations were unsurprisingly picked up by more than one imputation approach, and thus we ultimately identified 92 unique gene-trait pairings. We identified significant associations for 15/29 traits (52%), with the most associations observed for SHBG (19), total bilirubin (11), apolipoprotein B (9), and lipoprotein A (9). In order to evaluate the utility of our LA-aware approaches, we compare and contrast the total number of unique gene-trait associations identified in Figure S16. Leveraging our LA-derived ancestry-specific p-values (aspPS AFR/EUR) and the p-values from their combined component (casPS), this Level 2 aggregation successfully identified 83/92 associations (90.2%). 18 gene-trait associations were identified by this casPS+aspPS approach that were not identified by standard GReX imputation using reference eQTL summary data (PS AFR or PS EUR). Some of the genes among these 18 gene-trait pairs are located in close proximity to each other and likely do not reflect independent regions of association. Therefore, in Table 1, we present the most significant gene observed within a 1Mb window for each phenotype. We provide the full lists of all gene-trait associations identified by each individual approach in the supplement (Tables S1-S5).

**Table 1.** Associations identified in UKB blood biomarker analysis by Level 2 p-value aggregation of LA-aware casPS and aspPSs that were not identified using standard GReX imputation using PS trained in reference AFR and EUR eQTL summary data. For genomic loci (1MB regions) containing multiple identified genes for the same phenotype, only the most significant gene is presented. The p-values shown below represent ACAT(casPS p, aspPS AFR p, aspPS EUR p).

| Phenotype | Group | Gene | Gene Type | Chr | Pos (hg38) | p |
| --- | --- | --- | --- | --- | --- | --- |
| Direct bilirubin | Liver | *ATG16L1* | Protein coding | 2 | 233210051 | 3.11E-09 |
| Lipoprotein A | Cardiovascular | *ZDHHC14* | Protein coding | 6 | 157381133 | 6.09E-08 |
| Lipoprotein A | Cardiovascular | *HNRNPH1P1* | Processed pseudogene | 6 | 159712801 | 1.11E-10 |
| Apolipoprotein A | Cardiovascular | *CD36* | Protein coding | 7 | 80369575 | 7.14E-08 |
| Apolipoprotein B | Cardiovascular | *IST1* | Protein coding | 16 | 71845996 | 1.08E-07 |
| SHBG | Hormone | *WRAP53* | Protein coding | 17 | 7686071 | 3.35E-17 |
| Cholesterol | Cardiovascular | *ZNF491* | Protein coding | 19 | 11797667 | 7.96E-08 |
| LDL direct | Cardiovascular | *ZNF491* | Protein coding | 19 | 11797667 | 6.26E-09 |
| Gamma glutamyltransferase | Liver | *LRRC75B* | Protein coding | 22 | 24585620 | 5.39E-10 |
| Creatinine | Renal | *XRCC6* | Protein coding | 22 | 41621119 | 1.40E-09 |

Of the 9 genes identified by the casPS+aspPS approach highlighted in Table 1, the vast majority have consistent evidence in the GWAS literature, harboring one or more significant GWAS variant (p < 5E-08) for the relevant traits. However, one association (*HNRNPH* for lipoprotein A) is not located near known GWAS loci^67^. While *HNRNPH* is a pseudogene, and thus the biological mechanisms behind this association are thus unclear, previous work has helped elucidate the role that pseudogenes can play in the development of cardiovascular disease^68^.

Next, we used the online database TWAS Atlas to examine prior documented TWAS associations for each of these 9 genes^3^. Some of the genes identified exclusively by the LA-aware approach have been implicated in previous TWAS of relevant traits, while others represent potentially new TWAS findings. For example, we did not find prior TWAS associations of *ATG16L1* with traits relevant to bilirubin levels. While some of the genes identified by our approach for lipoproteins have been implicated by previous TWAS of lipid biomarkers of cardiovascular disease (e.g., *ZDHHC14, CD36, IST1*), *HNRNPH1P1* does not have any reported association in the database. The gene associated with SHBG in Table 1 (*WRAP53*) was identified in association with endometriosis in a recent TWAS^69^. SHBG plays a role in the availability of sex hormones in the body, and increased levels of SHBG were observed among women with endometriosis compared to controls^70^.While *ZNF491* has been identified by previous TWASs of cholesterol, genes *LRRC75B* and *XRCC6* did not have any significant prior TWAS associations with biomarker-relevant phenotypes in the Atlas.

Lastly, we note that the genes highlighted above were identified by Level 2 p-value aggregation of casPS and the ancestry-specific partial GReX results (casPS+aspPS). We focus on these findings to underscore the additional insight that can be found by LA-aware approaches only. However, CADET optimally allows for further aggregation of results from the LA-unaware standard GReX imputation approaches (i.e., casPS+aspPS+PS). Using this method, we identify four additional significant gene-trait associations (Figure S16, Table S2).

## 4. DISCUSSION

In this project, we introduce CADET, a method for performing transcriptome-wide association analysis in admixed subjects. Genomes of admixed individuals are a mosaic of two or more distinct ancestral groups, and thus local ancestry tract information within each haplotype can be leveraged to improve power in genetic association analyses when causal variant effect sizes differ between populations. Our method contributes to the growing catalog of statistical methods dedicated to admixed individuals with the appealing feature that it does not require individual-level genotype and gene expression training data, as is typically needed in Stage 1 GReX model training in TWASs. Here, we build upon a recently published TWAS approach, OTTERS, that uses well-established PRS models designed for GWAS summary data and applies them to eQTL summary data to impute gene expression in a tissue of interest^16^. In CADET, we use multiple sets of eQTL summary statistics, namely those derived from the distinct parent ancestral groups of our admixed testing sample. The framework is flexible enough to also incorporate eQTL summary data from admixed cohorts independent of the testing set. The CADET software is available on GitHub (Web Resources).

Through our simulations, we first demonstrate that p-values of all variants of our proposed approach yield the expected distribution under the null assumption of no gene-trait association. These variants include two approaches of p-value aggregation: Level (1), wherein we combine p-values from the three PRS models, and Level (2), in which we aggregate Level (1) p-values. Next, through our power simulations, we make several important observations. First, under all scenarios of 10 or fewer eQTLs, both our Level (1) casPS and Level (2) (casPS+aspPS+PS) aggregation methods achieve superior power to any standard LA-unaware approach when the genetic architecture of gene expression differs between AFR and EUR populations. In fact, there is growing evidence for such ancestry-specific genetic architecture patterns, and estimated gene expression heritability has also been shown to differ significantly by local ancestry at the transcription start site of genes among admixed subjects^17^. Second, even when genetic architecture patterns are identical between ancestries, our LA-aware Level (1) casPS and Level (2) (casPS+aspPS+PS) generally achieve greater power or power comparable to if we employed standard GReX imputation using eQTL summary data from a perfectly matched (number of admixture generations, initial AFR/EUR contribution proportions) independent admixed sample. Third, we also see that our Level (2) approach still performs competitively in these settings when we assume a large number (10%) of local ancestries are misclassified in the gene region and is the preferred method for use in practice.

Finally, in our applied analysis, we use real-world eQTL summary data from a European sample and a high-African ancestry sample of African Americans to perform TWAS of 29 blood biomarker traits. Here, our testing set is two-way African and European admixed subjects from the UK Biobank. We successfully identified 18 significant gene-trait associations using our Level 2 p-value aggregation approach (casPS + aspPSs) that were not picked up using standard GReX PS imputation methods that ignore local ancestry (PS AFR, PS EUR). The majority of these genes are consistent with GWAS loci previously identified for the corresponding traits. However, one gene, *HNRNPH* on chromosome 6, represents a potentially novel locus for lipoprotein A and was not identified by standard GReX imputation.

We observe several limitations of our present work. We first note that while CADET demonstrates desirable power levels for modest trait heritability, the gene expression imputation accuracies are lower across all simulation settings than the simulated gene expression heritability levels. We argue that our imputation models are trained using PRS approaches to leverage the widespread availability of eQTL summary data, and R^2^ has been shown to fall below true trait heritability for common PRS approaches^16,65^, and we expect imputation accuracy to be even lower when ancestry-specific effects are involved. We argue that we may further improve imputation accuracy in CADET by including more PRS approaches to our method, and, importantly, those designed precisely for admixed samples (e.g., GAUDI^42^). We also note that our framework is flexible enough to incorporate individual-level training data (paired genotype and gene expression information), if available. However, previous work has shown that GReX models trained with individual-level reference data showed modest to no improvement in testing R^2^ over various PRS-based models trained with summary data^16^. Additionally, in this project, we only consider two-way admixed individuals for both our simulated analyses and applied work in the UK Biobank. We believe that we can easily extend this approach to allow for three-way or higher levels of admixture, and that the approach is computationally efficient enough to implement this in practice. Finally, we designed our method to utilize eQTL data from ancestrally homogeneous parent ancestral groups. However, when seeking to apply our method to admixed individuals of African ancestry, we note there is a marked paucity of eQTL studies performed in non-admixed African cohorts. The eQTL summary data from African American individuals with >50% African ancestry-alleles currently represents the best surrogate dataset of reasonable sample size for our analysis, and this substitution has similarly been employed in another recent methodological paper^42^.

## Acknowledgements

This work was supported by the National Institutes of Health [AG071170, CA211574].

## Web Resources

The code for implementing this method is publicly available at https://github.com/staylorhead/CADET.

## Data and Code Availability

The GTEx V8 cis-eQTL summary statistics in Europeans used for the applied analysis of UKB subjects are publicly available and can be accessed at https://www.gtexportal.org/home/downloads/adult-gtex/qtl. The cis-eQTL summary statistics from high-African ancestry African Americans by Kachuri et al.^17^ also used in our applied UKB analysis can be accessed at https://zenodo.org/records/7735723.

