## Supplemental Figures for "CADET: Enhanced transcriptome-wide association analyses in admixed samples using eQTL summary data"

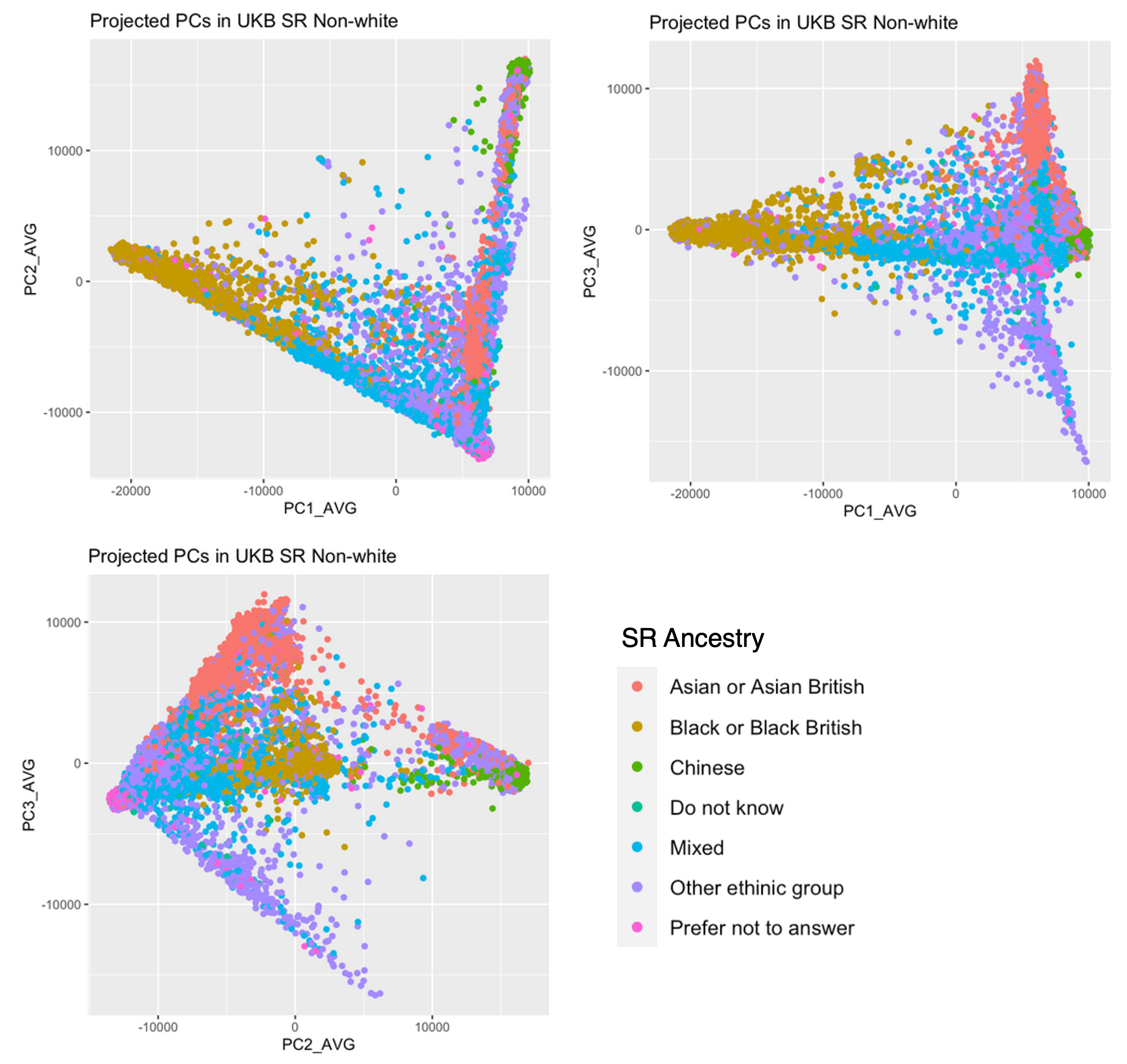

**Figure S1.** Projection of UKB self-reported (SR) non-White individuals (N=27,491) onto three-dimensional principal component space calculated using 1000 Genomes samples from the following superpopulations: African, American, East Asian, European and South Asian. Coloring of samples indicates self-reported ethnicity of UKB subjects.

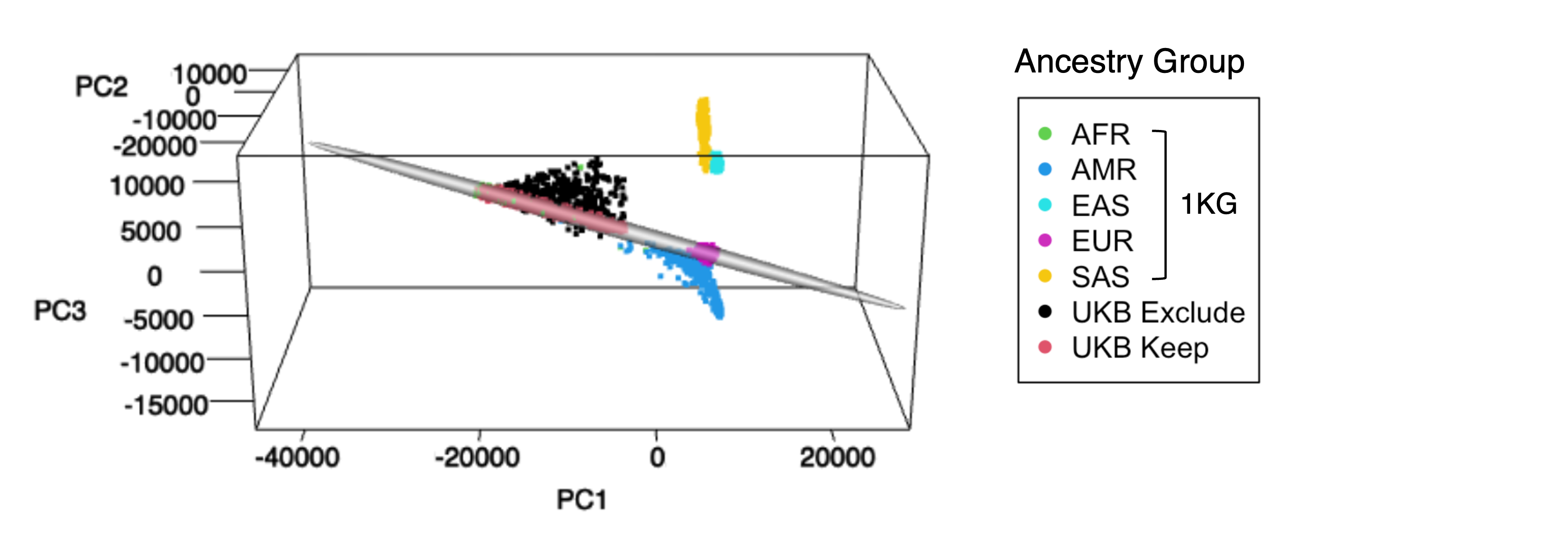

**Figure S2.** Projection onto 1000 Genomes principal component space of N=9,187 UKB self-reported non-white subjects with >50% probability of AFR ancestry by the random forest ancestry classification model. These axes were calculated using reference samples from the following superpopulations: African (AFR), American (AMR), East Asian (EAS), European (EUR), and South Asian (SAS). These subjects are also included on the plot to provide orientation to UKB subjects. UKB subjects (red, black) are colored by whether they fall within the 95% ellipsoid along the AFR-EUR cline.

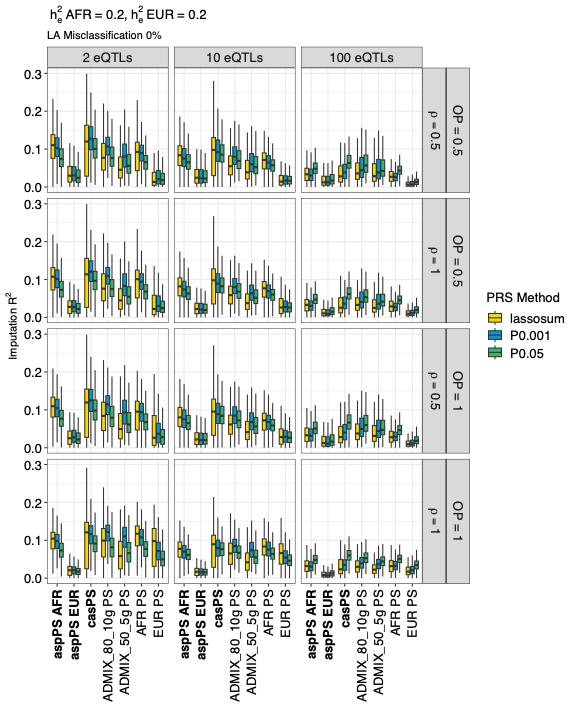

**Figure S3.** Gene expression imputation accuracy in 10,000 admixed testing samples (10 admixture generations, 80% initial contribution from AFR, 20% initial contribution from EUR) for expression heritability $h_{e,AFR}^{2},h_{e,EUR}^{2}=0.2$. Vertical panels indicate the true number of causal SNPs for gene expression (eQTLs). Horizontal panels indicate the proportion of eQTLs that overlap (OP) between AFR and EUR ancestries, as well as the correlation in eQTL effect sizes for shared eQTLs between the two ancestral groups (*ρ*). The x-axis shows the GReX imputation approach, including our proposed local-ancestry aware methods (aspPS, casPS, bold) and standard PRS imputation approaches (PS). For ancestry-aware methods, we assume no local ancestry misclassification. Whiskers of boxplot extend to maximum/minimum point that is less than 1.5*IQR from the third/first quartiles.

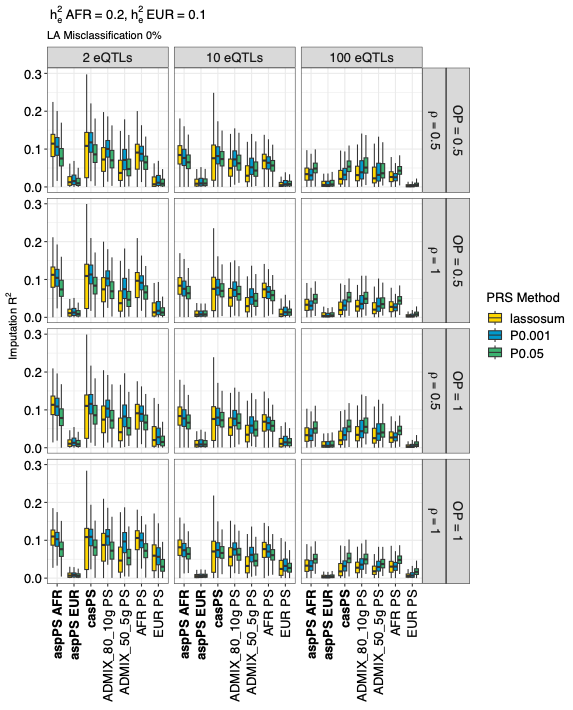

**Figure S4.** Gene expression imputation accuracy in 10,000 admixed testing samples (10 admixture generations, 80% initial contribution from AFR) for expression heritability $h_{e,AFR}^{2}=0.2,h_{e,EUR}^{2}=0.1$. Vertical panels indicate the true number of causal SNPs for gene expression (eQTLs). Horizontal panels indicate the proportion of eQTLs that overlap (OP) between AFR and EUR ancestries, as well as the correlation in eQTL effect sizes for shared eQTLs between the two ancestral groups (*ρ*). The x-axis shows the GReX imputation approach, including our proposed local-ancestry aware methods (aspPS, casPS, bold) and standard PRS imputation approaches (PS). For ancestry-aware methods, we assume no local ancestry misclassification. Whiskers of boxplot extend to maximum/minimum point that is less than 1.5*IQR from the third/first quartiles.

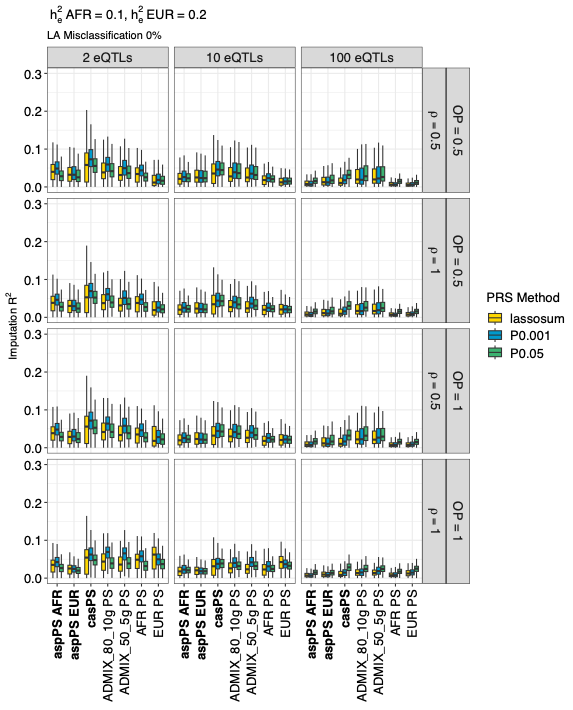

**Figure S5.** Gene expression imputation accuracy in 10,000 admixed testing samples (10 admixture generations, 80% initial contribution from AFR) for expression heritability $h_{e,AFR}^{2}=0.1,h_{e,EUR}^{2}=0.2$. Vertical panels indicate the true number of causal SNPs for gene expression (eQTLs). Horizontal panels indicate the proportion of eQTLs that overlap (OP) between AFR and EUR ancestries, as well as the correlation in eQTL effect sizes for shared eQTLs between the two ancestral groups (*ρ*). The x-axis shows the GReX imputation approach, including our proposed local-ancestry aware methods (aspPS, casPS, bold) and standard PRS imputation approaches (PS). For ancestry-aware methods, we assume no local ancestry misclassification. Whiskers of boxplot extend to maximum/minimum point that is less than 1.5*IQR from the third/first quartiles.

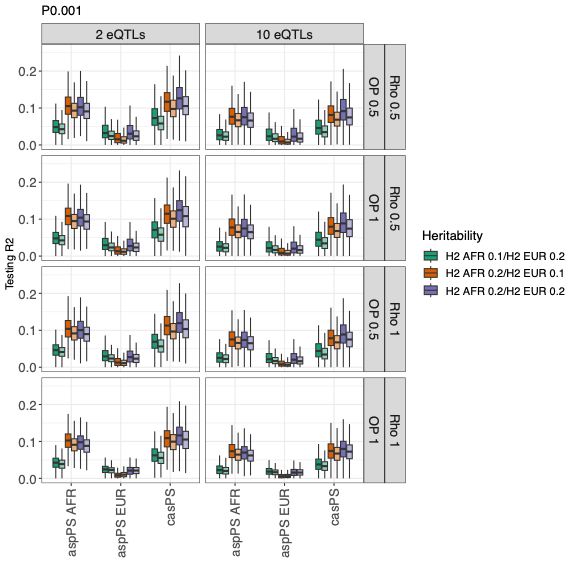

**Figure S6**. Gene expression imputation accuracy of pruning and thresholding (PT = 0.001) for LA-aware approaches in 10,000 admixed testing samples (10 admixture generations, 80% initial contribution from AFR). Vertical panels indicate the true number of causal SNPs for gene expression (eQTLs). Horizontal panels indicate the proportion of eQTLs that overlap (OP) between AFR and EUR ancestries, as well as the correlation in eQTL effect sizes for shared eQTLs between the two ancestral groups (ρ). For these ancestry-aware methods, we assume either no LA misclassification (darker boxes) or 10% misclassification of SNPs in the gene region (lighter boxes). Whiskers of boxplot extend to maximum/minimum point that is less than 1.5*IQR from the third/first quartiles.

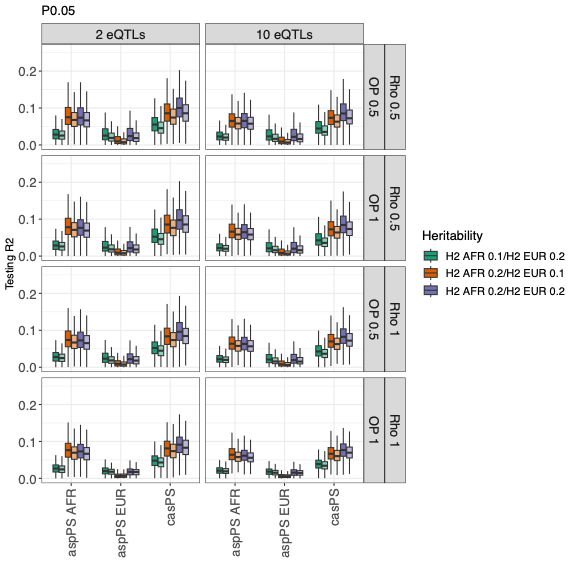

**Figure S7**. Gene expression imputation accuracy of pruning and thresholding (PT = 0.05) for LA-aware approaches in 10,000 admixed testing samples (10 admixture generations, 80% initial contribution from AFR). Vertical panels indicate the true number of causal SNPs for gene expression (eQTLs). Horizontal panels indicate the proportion of eQTLs that overlap (OP) between AFR and EUR ancestries, as well as the correlation in eQTL effect sizes for shared eQTLs between the two ancestral groups (ρ). For these ancestry-aware methods, we assume either no LA misclassification (darker boxes) or 10% misclassification of SNPs in the gene region (lighter boxes). Whiskers of boxplot extend to maximum/minimum point that is less than 1.5*IQR from the third/first quartiles.

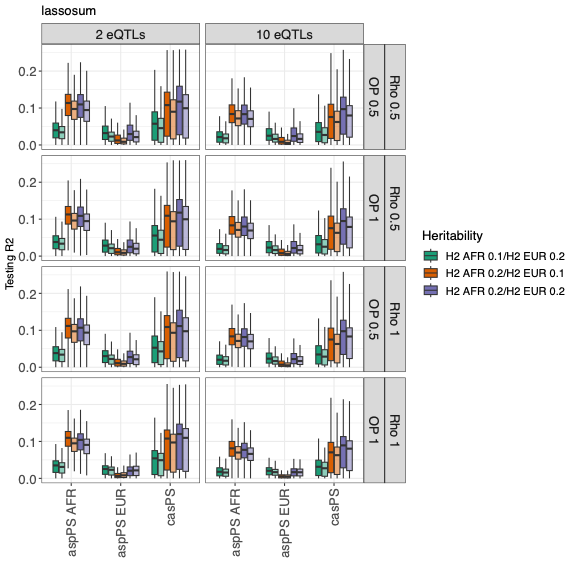

**Figure S8**. Gene expression imputation accuracy of lassosum for LA-aware approaches in 10,000 admixed testing samples (10 admixture generations, 80% initial contribution from AFR). Vertical panels indicate the true number of causal SNPs for gene expression (eQTLs). Horizontal panels indicate the proportion of eQTLs that overlap (OP) between AFR and EUR ancestries, as well as the correlation in eQTL effect sizes for shared eQTLs between the two ancestral groups (ρ). For these ancestry-aware methods, we assume either no LA misclassification (darker boxes) or 10% misclassification of SNPs in the gene region (lighter boxes). Whiskers of boxplot extend to maximum/minimum point that is less than 1.5*IQR from the third/first quartiles.

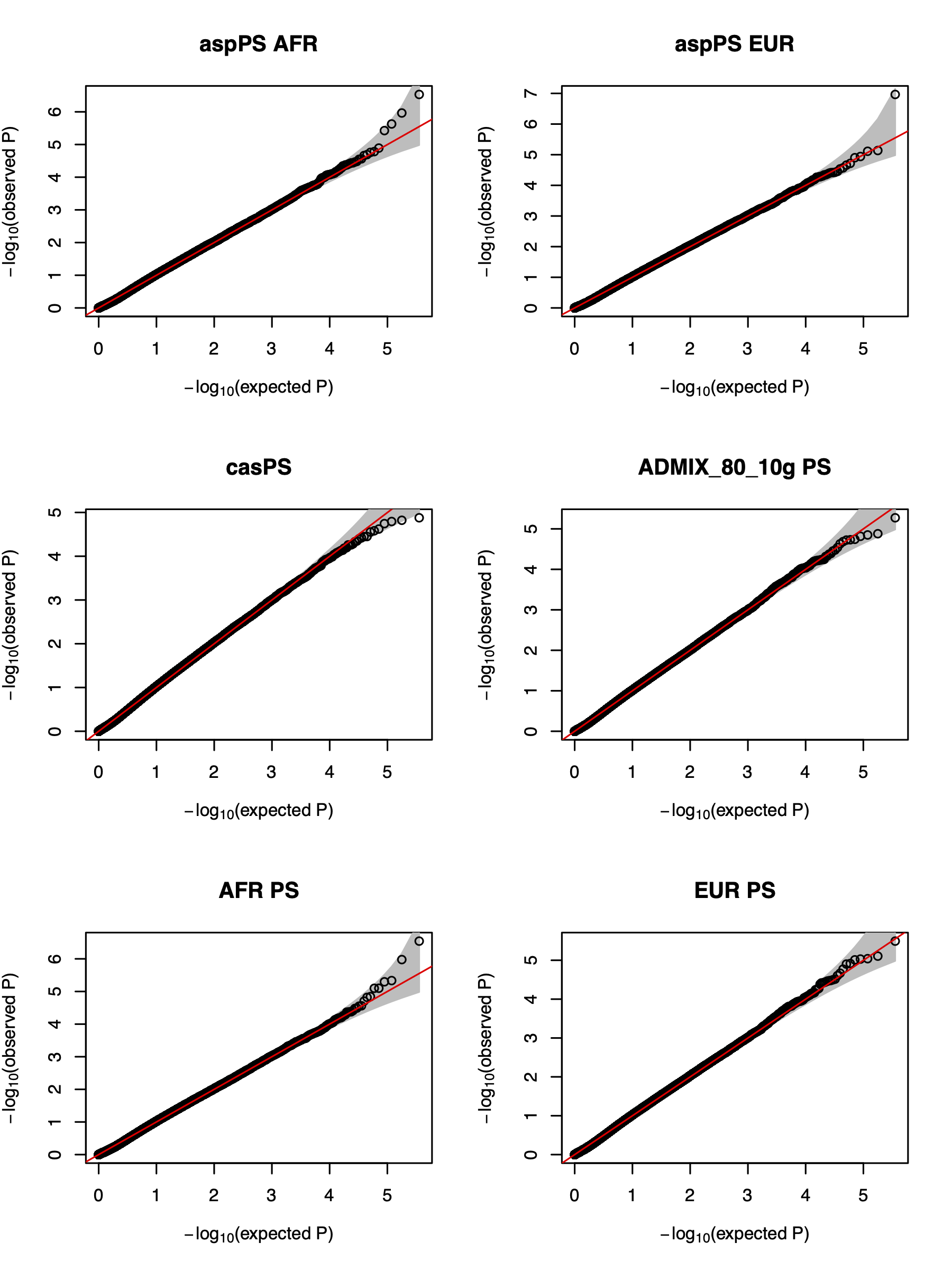

**Figure S9.** QQ plots of p-values from gene-level association tests from both LA-aware and LA-unaware GReX imputation approaches under the null when no association of expression with trait exists. Here, we assume a testing sample size of 10,000. These p-values represent Level 1 p-value aggregation by ACAT, i.e., aggregation of p-values across the three PRS models (P+T0.001, P+T0.05, lassosum). For ancestry-aware methods, we assume no local ancestry misclassification. Each plot shown corresponds to 36,000 total simulations, including all 36 gene expression simulation settings.

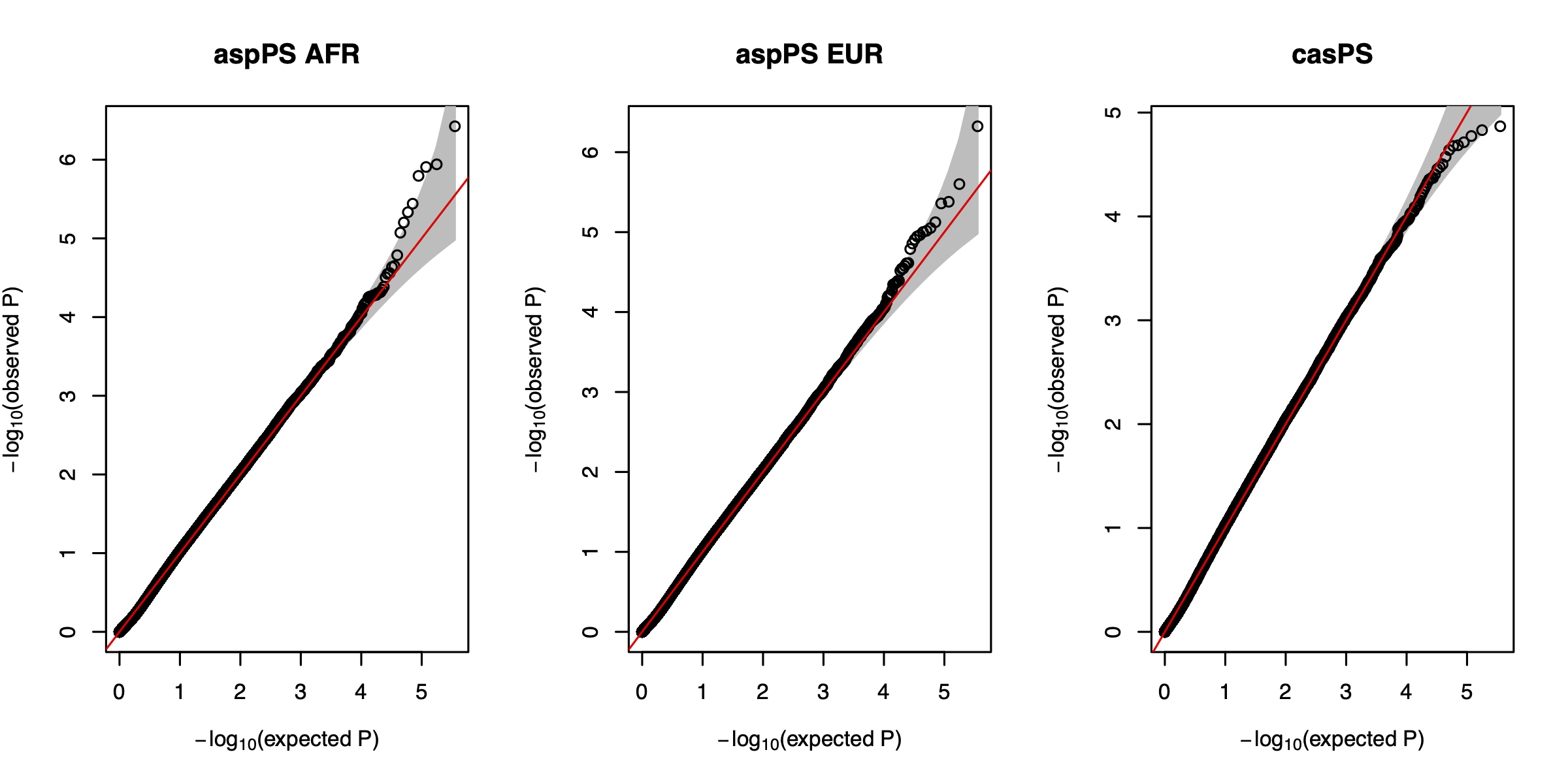

**Figure S10**. QQ plots of p-values from gene-level association tests from LA-aware GReX imputation approaches under the null when no association of expression with trait exists. For these ancestry-aware methods, we assume 10% local ancestry mis- classification for SNPs in the gene region. Here, we assume a testing sample size of 10,000. These p-values represent Level 1 p-value aggregation by ACAT, i.e., aggregation of p-values across the three PRS models (P+T0.001, P+T0.05, lassosum). Each plot shown corresponds to 36,000 total simulations, including all 36 gene expression simulation settings.

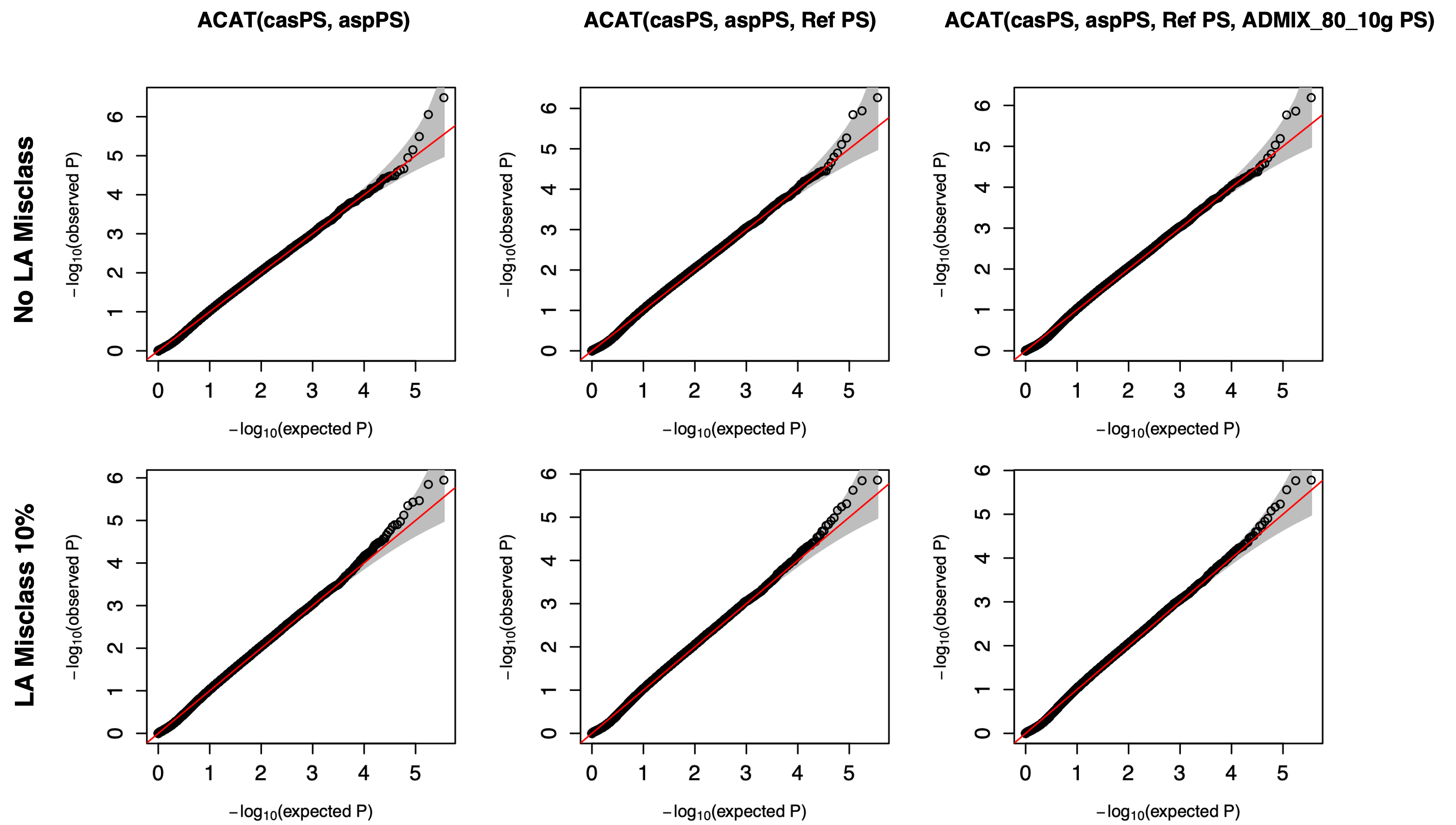

**Figure S11.** QQ plots of p-values from gene-level association tests from LA-aware GReX imputation approaches under the null when no association of expression with trait exists. Here, we assume a testing sample size of 10,000. These p-values represent Level 2 p-value aggregation by ACAT. The p-value aggregation approach is indicated in the plot title. For ancestry-aware methods, we assume either no local ancestry misclassification (No LA Misclass) or misclassification of 10% of SNPs in the gene region (LA Misclass 10%). Each plot shown corresponds to 36,000 total simulations, including all 36 gene expression simulation settings.

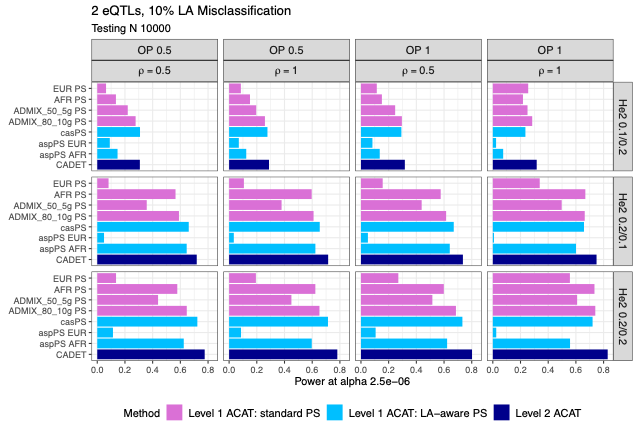

**Figure S12**. Power of gene-level association tests of imputed GReX vectors and simulated trait at significance level 2.5E-6. Here, we assume a phenotypic heritability of $h_{p}^{2}$ = 0.025, 2 eQTLs, 10% local ancestry (LA) misclassification for LA-aware approaches, and a testing dataset sample size of 10,000. Vertical panels indicate the proportion of eQTLs that are shared between AFR and EUR ancestries (OP) and the correlation of eQTL effect sizes for shared eQTLs ($\rho$). Horizontal panels indicate the gene expression heritability in AFR and EUR ancestries ($h_{e}^{2}$AFR/EUR). Pink bars indicate the power of LA-unaware GReX imputation approaches, with p-values aggregated across the three PRS models (ACAT Level 1). Light blue bars indicate LA-aware approaches with Level 1 p-value aggregation by ACAT. Dark blue bars indicate the power of LA-aware approaches, aggregating both PRS p-values and the resulting p-values of casPS, aspPSs (AFR and EUR), and standard PSs trained in the two AFR/EUR reference populations (ACAT Level 2).

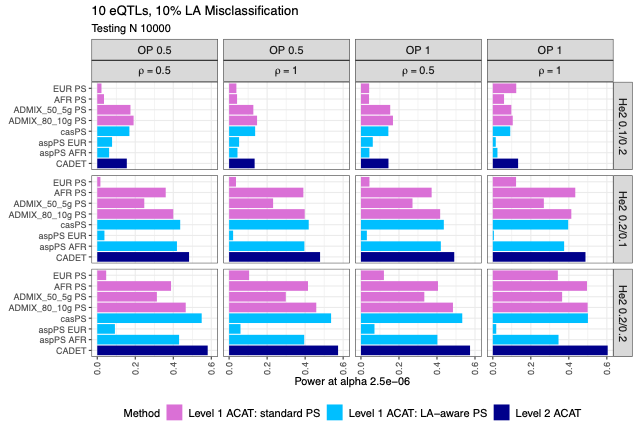

**Figure S13**. Power of gene-level association tests of imputed GReX vectors and simulated trait at significance level 2.5E-6. Here, we assume a phenotypic heritability of $h_{p}^{2}$ = 0.025, 10 eQTLs, 10% local ancestry (LA) misclassification for LA-aware approaches, and a testing dataset sample size of 10,000. Vertical panels indicate the proportion of eQTLs that are shared between AFR and EUR ancestries (OP) and the correlation of eQTL effect sizes for shared eQTLs ($\rho$). Horizontal panels indicate the gene expression heritability in AFR and EUR ancestries ($h_{e}^{2}$AFR/EUR). Pink bars indicate the power of LA-unaware GReX imputation approaches, with p-values aggregated across the three PRS models (ACAT Level 1). Light blue bars indicate LA-aware approaches with Level 1 p-value aggregation by ACAT. Dark blue bars indicate the power of LA-aware approaches, aggregating both PRS p-values and the resulting p-values of casPS, aspPSs (AFR and EUR), and standard PSs trained in the two AFR/EUR reference populations (ACAT Level 2).

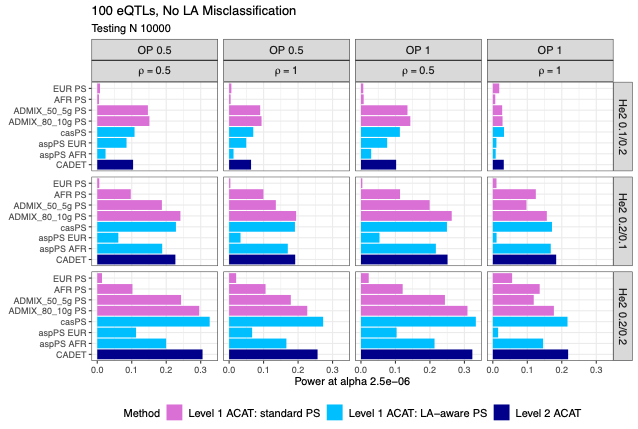

**Figure S14.** Power of gene-level association tests of imputed GReX vectors and simulated trait at significance level 2.5E-6. Here, we assume a phenotypic heritability of $h_{p}^{2}$ = 0.025, 100 eQTLs, no local ancestry (LA) misclassification for LA-aware approaches, and a testing dataset sample size of 10,000. Vertical panels indicate the proportion of eQTLs that are shared between AFR and EUR ancestries (OP) and the correlation of eQTL effect sizes for shared eQTLs ($\rho$). Horizontal panels indicate the gene expression heritability in AFR and EUR ancestries ($h_{e}^{2}$AFR/EUR). Pink bars indicate the power of LA-unaware GReX imputation approaches, with p-values aggregated across the three PRS models (ACAT Level 1). Light blue bars indicate LA-aware approaches with Level 1 p-value aggregation by ACAT. Dark blue bars indicate the power of LA-aware approaches, aggregating both PRS p-values and the resulting p-values of casPS, aspPSs (AFR and EUR), and standard PSs trained in the two AFR/EUR reference populations (ACAT Level 2).

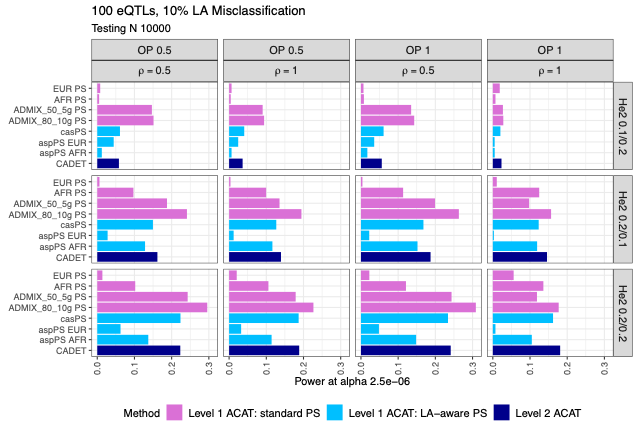

**Figure S15.** Power of gene-level association tests of imputed GReX vectors and simulated trait at significance level 2.5E-6. Here, we assume a phenotypic heritability of $h_{p}^{2}$ = 0.025, 100 eQTLs, 10% local ancestry (LA) misclassification for LA-aware approaches, and a testing dataset sample size of 10,000. Vertical panels indicate the proportion of eQTLs that are shared between AFR and EUR ancestries (OP) and the correlation of eQTL effect sizes for shared eQTLs ($\rho$). Horizontal panels indicate the gene expression heritability in AFR and EUR ancestries ($h_{e}^{2}$AFR/EUR). Pink bars indicate the power of LA-unaware GReX imputation approaches, with p-values aggregated across the three PRS models (ACAT Level 1). Light blue bars indicate LA-aware approaches with Level 1 p-value aggregation by ACAT. Dark blue bars indicate the power of LA-aware approaches, aggregating both PRS p-values and the resulting p-values of casPS, aspPSs (AFR and EUR), and standard PSs trained in the two AFR/EUR reference populations (ACAT Level 2).

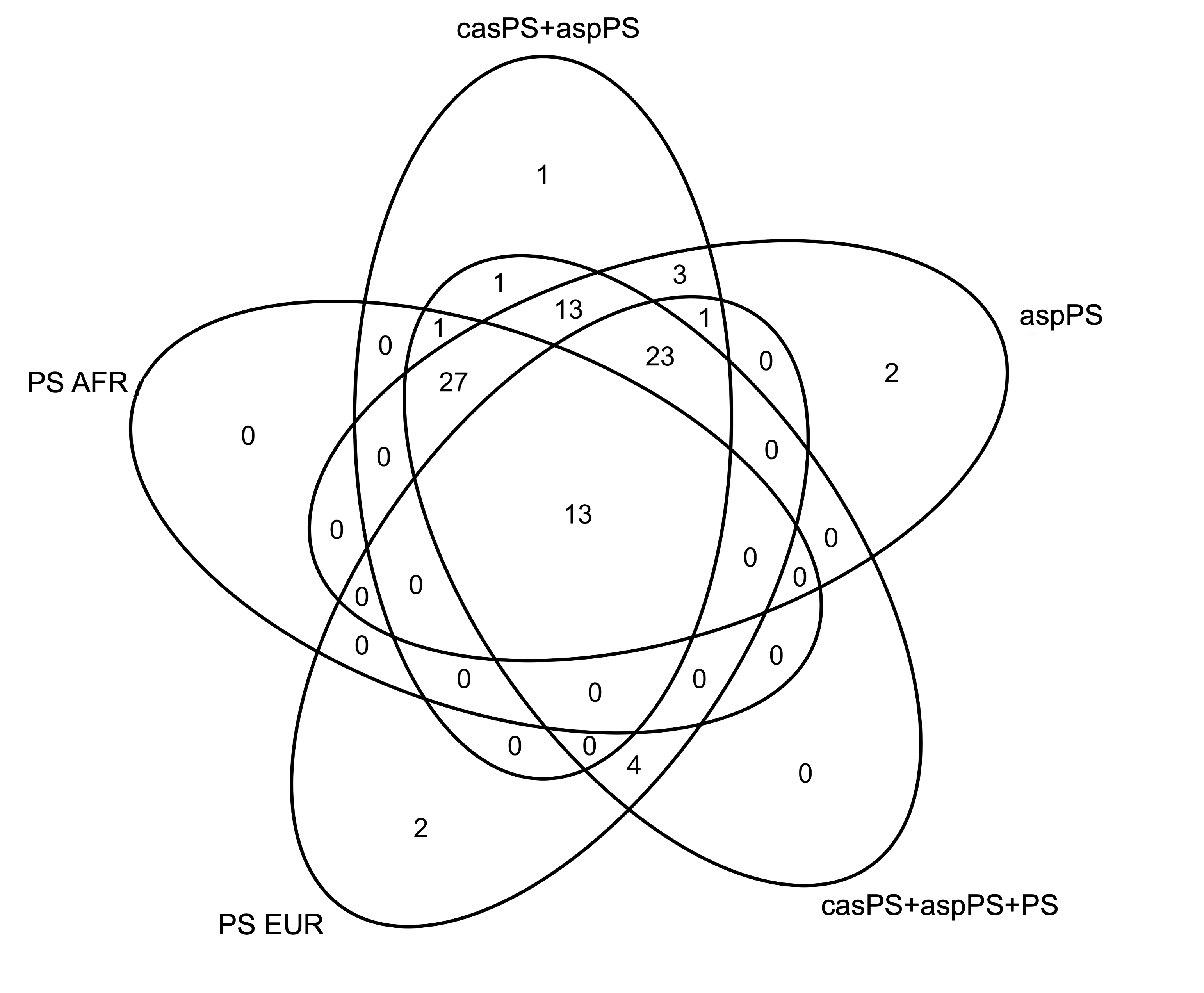

**Figure S16**. Number of significant gene-trait associations found across all 29 blood biomarker traits in UKB analysis. Counts are grouped by GReX imputation and p-value aggregation approach. We see 18 significant gene-trait associations identified for casPS+aspPS (Level 2 ACAT aggregation approach of only LA-aware PSs) that were not found by PS AFR or PS EUR, as these are indicative of hits that are found only using LA-aware methods and utilize no p-values from standard PS approaches. The final CADET results (casPS+aspPS+PS) combine p-values from both LA-unaware and LA-aware GReX imputation approaches.
