## Supplemental Information for "CADET: Enhanced transcriptome-wide association analyses in admixed samples using eQTL summary data"

### Derivation of Gene Expression Heritability in Admixed Individuals

Let there be $i = 1, ..., N$ admixed individuals in our testing dataset. Assume there are $V$ total causal SNPs in each ancestry for this phenotype (e.g., gene expression). The $V$ causal SNPs for EUR are *not* assumed to be the same as the $V$ causal SNPs for AFR.

In general, subscript 1 corresponds to AFR and subscript 2 corresponds to EUR.

- $x_{ivm1} \in \{0, 1\}$ : number of minor alleles for $v$th causal AFR SNP on maternal haplotype of subject $i$
- $x_{ivp1} \in \{0, 1\}$ : number of minor alleles for $v$th causal AFR SNP on paternal haplotype of subject $i$
- $x_{ivm2} \in \{0, 1\}$ : number of minor alleles for $v$th causal EUR SNP on maternal haplotype of subject $i$
- $x_{ivp2} \in \{0, 1\}$ : number of minor alleles for $v$th causal EUR SNP on paternal haplotype of subject $i$
- $\gamma_{ivm1} \in \{0, 1\}$ : 1 if AFR local ancestry of $v$th causal AFR SNP on maternal haplotype of subject $i$
- $\gamma_{ivp1} \in \{0, 1\}$ : 1 if AFR local ancestry of $v$th causal AFR SNP on paternal haplotype of subject $i$
- $\gamma_{ivm2} \in \{0, 1\}$ : 1 if EUR local ancestry of $v$th causal EUR SNP on maternal haplotype of subject $i$
- $\gamma_{ivp2} \in \{0, 1\}$ : 1 if EUR local ancestry of $v$th causal EUR SNP on paternal haplotype of subject $i$

Let $g_{iv1}$ represent the number of AFR-ancestry minor alleles of the $v$th causal AFR SNP and $g_{iv2}$ be the number of EUR-ancestry minor alleles of subject $i$ at $v$th causal EUR SNP:

$$g_{iv1} := x_{ivm1}\gamma_{ivm1} + x_{ivp1}\gamma_{ivp1} \tag{1}$$

$$g_{iv2} := x_{ivm2}\gamma_{ivm2} + x_{ivp2}\gamma_{ivp2} \tag{2}$$

We can arrange these into the following matrices, where each column has been centered:

$$G_1 = \begin{pmatrix} \underset{\sim}{g}_{11} & \cdots & \underset{\sim}{g}_{V1} \end{pmatrix}_{N \times V} \text{ where } \underset{\sim}{g}_{v1} = \begin{pmatrix} g_{1v1} - \bar{g}_{v1} \\ \vdots \\ g_{Nv1} - \bar{g}_{v1} \end{pmatrix}_{N \times 1} \text{ and } \bar{g}_{v1} = \frac{1}{N}\sum_{i=1}^{N} g_{iv1}$$

$$G_2 = \begin{pmatrix} \underset{\sim}{g}_{12} & \cdots & \underset{\sim}{g}_{V2} \end{pmatrix}_{N \times V} \text{ where } \underset{\sim}{g}_{v2} = \begin{pmatrix} g_{1v2} - \bar{g}_{v2} \\ \vdots \\ g_{Nv2} - \bar{g}_{v2} \end{pmatrix}_{N \times 1} \text{ and } \bar{g}_{v2} = \frac{1}{N}\sum_{i=1}^{N} g_{iv2}$$

The following quantities will be needed to derive heritability:

$$f_{v1} = \frac{\sum_{i=1}^{N} g_{iv1}}{\sum_{i=1}^{N} \gamma_{ivm1} + \sum_{i=1}^{N} \gamma_{ivp1}} = \text{African-specific MAF at } v\text{th causal AFR SNP}$$

$$f_{v2} = \frac{\sum_{i=1}^{N} g_{iv2}}{\sum_{i=1}^{N} \gamma_{ivm2} + \sum_{i=1}^{N} \gamma_{ivp2}} = \text{European-specific MAF at } v\text{th causal EUR SNP}$$

$\theta_{v1} = \frac{\sum_{i=1}^{N}\gamma_{ivm1}+\sum_{i=1}^{N}\gamma_{ivp1}}{2N}$ = proportion of alleles at $v$th causal AFR SNP that are African ancestry

$\theta_{v2} = \frac{\sum_{i=1}^{N}\gamma_{ivm2}+\sum_{i=1}^{N}\gamma_{ivp2}}{2N}$ = proportion of alleles at $v$th causal EUR SNP that are European ancestry

$\theta_{v12} = \theta_{v1}\left(\frac{\sum_{i=1}^{N}\gamma_{ivm1}(\gamma_{ivm2})+\sum_{i=1}^{N}\gamma_{ivp1}(\gamma_{ivp2})}{\sum_{i=1}^{N}(\gamma_{ivm2})+\sum_{i=1}^{N}(\gamma_{ivp2})}\right)$ = probability that $v$th causal AFR SNP is African ancestry and $v$th causal EUR SNP is European ancestry (don't actually end up using this quantity)

These definitions imply:

$\sum_{i=1}^{N} g_{iv1} = 2N\theta_{v1}f_{v1}$

$\sum_{i=1}^{N} g_{iv2} = 2N\theta_{v2}f_{v2}$

$\bar{g}_{v1} = 2\theta_{v1}f_{v1}$

$\bar{g}_{v2} = 2\theta_{v2}f_{v2}$

Let us assume that we want to standardize genotypes by local ancestry.

We also assume that, of the $V$ causal SNPs in each ancestry, the first $S$ are shared between AFR and EUR, and the remaining $U$ are unique to each ancestry ($S+U=V$). We can therefore model the $N\times 1$ phenotype outcome vector (gene expression) $\underset{\sim}{y}$ as:

$\underset{\sim}{y} = G_1T_1^{1/2}\dot{\beta}_1 + G_2T_2^{1/2}\dot{\beta}_2 + \underset{\sim}{\epsilon}$

Here, $\underset{\sim}{\epsilon} \sim N(0,(1-h^2_{adm})I_N)$ and $\dot{\beta}_1, \dot{\beta}_2$ represent the $V\times 1$ vectors of ancestry-specific effects per genotype standard deviation. The phenotypic heritability is $h^2_{adm}$.

$T_1$ is a $V\times V$ diagonal matrix with $(T_1)_{vv} = \tau^2_{v1} = \frac{1}{2f_{v1}(1-f_{v1})}$

$T_2$ is a $V\times V$ diagonal matrix with $(T_2)_{vv} = \tau^2_{v2} = \frac{1}{2f_{v2}(1-f_{v2})}$

Let $h_1^2$ be heritability of phenotype in AFR ancestry and $h_2^2$ be heritability of phenotype in EUR ancestry.

Let $\rho = \text{Corr}(\dot{\beta}_{1v}, \dot{\beta}_{2v}), v \in \{1,...,S\}$, i.e., the correlation of ancestry-specific effects for causal SNPs that are common to both ancestries.

$$\begin{pmatrix}\dot{\beta}_1\\ \dot{\beta}_2\end{pmatrix}_{2V\times 1} = \begin{pmatrix}\dot{\beta}_{1S}\\ \dot{\beta}_{1U}\\ \dot{\beta}_{2S}\\ \dot{\beta}_{2U}\end{pmatrix} \sim N\left(\begin{pmatrix}0\\ \vdots\\ \vdots\\ 0\end{pmatrix}, \begin{pmatrix}\frac{h_1^2}{V}I_S & 0_{S\times U} & \frac{\rho}{V}\sqrt{h_1^2h_2^2}I_S & 0_{S\times U}\\ & \frac{h_1^2}{V}I_U & 0_{U\times S} & 0_{U\times U}\\ & & \frac{h_2^2}{V}I_S & 0_{S\times U}\\ & & & \frac{h_2^2}{V}I_U\end{pmatrix}\right)$$

To derive the heritability in admixed subjects, let's rewrite $G_1T_1^{1/2}\dot{\beta}_1$ as $G_1\beta_1$ and $G_2T_2^{1/2}\dot{\beta}_2$ as $G_2\beta_2$, where:

$$\begin{pmatrix}\beta_1\\ \beta_2\end{pmatrix}_{2V\times 1} = \begin{pmatrix}\beta_{1S}\\ \beta_{1U}\\ \beta_{2S}\\ \beta_{2U}\end{pmatrix} \sim N\left(\begin{pmatrix}0\\ \vdots\\ \vdots\\ 0\end{pmatrix}, \begin{pmatrix}A & 0_{S\times U} & B & 0_{S\times U}\\ & C & 0_{U\times S} & 0_{U\times U}\\ & & D & 0_{S\times U}\\ & & & E\end{pmatrix}\right)$$

$A_{S\times S} = \text{Cov}(\beta_{1S}, \beta_{1S}) = \frac{h_1^2}{V}diag(\tau^2_{v1}), v = 1,...,S$

$B_{S\times S} = \text{Cov}(\beta_{1S}, \beta_{2S}) = \frac{\rho}{V}\sqrt{h_1^2h_2^2}diag(\tau_{v1}\tau_{v2}), v = 1,...,S$

$C_{U\times U} = \text{Cov}(\beta_{1U}, \beta_{1U}) = \frac{h_1^2}{V} diag(\tau_{v1}^2), v = S+1, ..., V$

$D_{S\times S} = \text{Cov}(\beta_{2S}, \beta_{2S}) = \frac{h_2^2}{V} diag(\tau_{v2}^2), v = 1, ..., S$

$E_{U\times U} = \text{Cov}(\beta_{2U}, \beta_{2U}) = \frac{h_2^2}{V} diag(\tau_{v2}^2), v = S+1, ..., V$

Since the variance of the phenotype is 1, we can define $h_{adm}^2 = \text{Var}(G_1\beta_1 + G_2\beta_2)$.

$\text{Var}(G_1\beta_1 + G_2\beta_2) =? = \frac{1}{N} tr(E[(G_1\beta_1 + G_2\beta_2)(G_1\beta_1 + G_2\beta_2)'])$

$= \frac{1}{N} tr(E[G_1\beta_1\beta_1'G_1' + G_2\beta_2\beta_1'G_1' + G_1\beta_1\beta_2'G_2' + G_2\beta_2\beta_2'G_2'])$

$= \frac{1}{N} tr(E[G_1\beta_1\beta_1'G_1'] + E[G_2\beta_2\beta_1'G_1'] + E[G_1\beta_1\beta_2'G_2'] + E[G_2\beta_2\beta_2'G_2'])$

$= \frac{1}{N}\{tr(E[G_1\beta_1\beta_1'G_1']) + tr(E[G_2\beta_2\beta_1'G_1']) + tr(E[G_1\beta_1\beta_2'G_2']) + tr(E[G_2\beta_2\beta_2'G_2'])\}$

We can rewrite each component in the brackets as follows:

$tr(E[G_1\beta_1\beta_1'G_1']) = E(tr[G_1\beta_1\beta_1'G_1'])$

$= E(tr[\beta_1\beta_1'G_1'G_1]$

$= tr(E[\beta_1\beta_1'G_1'G_1])$

$= tr[E(\beta_1\beta_1')G_1'G_1]$

$= tr(\text{Cov}[\beta_1, \beta_1]G_1'G_1)$

Since $\text{Cov}[\beta_1, \beta_1]$ is a diagonal matrix, the diagonal elements of the matrix product are the product of diagonal elements of each matrix.

First, let us derive some quantities needed to find the diagonal elements of $G'G$.

$\sum_{i=1}^{N} g_{iv1}^2 = \sum_{i=1}^{N} (x_{ivm1}\gamma_{ivm1} + x_{ivp1}\gamma_{ivp1})^2$

$= \sum_{i=1}^{N} (x_{ivm1}^2\gamma_{ivm1}^2 + 2x_{ivm1}x_{ivp1}\gamma_{ivm1}\gamma_{ivp1} + x_{ivp1}^2\gamma_{ivp1}^2)$

$= \sum_{i=1}^{N} x_{ivm1}\gamma_{ivm1} + 2\sum_{i=1}^{N} x_{ivm1}x_{ivp1}\gamma_{ivm1}\gamma_{ivp1} + \sum_{i=1}^{N} x_{ivp1}\gamma_{ivp1}$

$= N\bar{g}_{v1} + 2N\theta_{v1}^2 f_{v1}^2$

$= 2N\theta_{v1}f_{v1} + 2N\theta_{v1}^2 f_{v1}^2$

$= 2N\theta_{v1}f_{v1}(1 + \theta_{v1}f_{v1})$

Similarly, $\sum_{i=1}^{N} g_{iv2}^2 = 2N\theta_{v2}f_{v2}(1 + \theta_{v2}f_{v2})$

$\sum_{i=1}^{N} g_{iv1}g_{iv2} = \sum_{i=1}^{N} (x_{ivm1}\gamma_{ivm1} + x_{ivp1}\gamma_{ivp1})(x_{ivm2}\gamma_{ivm2} + x_{ivp2}\gamma_{ivp2})$

$= \sum_{i=1}^{N} x_{ivm1}\gamma_{ivm1}x_{ivm2}\gamma_{ivm2} + \sum_{i=1}^{N} x_{ivp1}\gamma_{ivp1}x_{ivm2}\gamma_{ivm2} + \sum_{i=1}^{N} x_{ivm1}\gamma_{ivm1}x_{ivp2}\gamma_{ivp2} + \sum_{i=1}^{N} x_{ivp1}\gamma_{ivp1}x_{ivp2}\gamma_{ivp2}$

If $v \leq S$, $\gamma_{ivm1} = 1 - \gamma_{ivm2}$ and $\theta_{v1} = 1 - \theta_{v2}$:

$\sum_{i=1}^{N} g_{iv1}g_{iv2} = \sum_{i=1}^{N} x_{ivp1}\gamma_{ivp1}x_{ivm2}\gamma_{ivm2} + \sum_{i=1}^{N} x_{ivm1}\gamma_{ivm1}x_{ivp2}\gamma_{ivp2}$

$= 2Nf_{v1}f_{v2}\theta_{v1}\theta_{v2}$

If $v > S$, the probability that the $v$th causal AFR SNP is of African ancestry and the probability that the $v$th causal EUR SNP is of European ancestry on the same haplotype are not independent, thus:

$\sum_{i=1}^{N} g_{iv1}g_{iv2} = 2Nf_{v1}f_{v2}\theta_{v1}\theta_{v2} + 2N\theta_{12}f_{v1}f_{v2}$

Now, to calculate the diagonal elements:

$(G_1'G_1)_{vv} = \underset{\sim}{g}_{v1}'\underset{\sim}{g}_{v1}$

$= \sum_{i=1}^{N}(g_{iv1} - \bar{g}_{v1})^2$

$= \sum_{i=1}^{N} g_{iv1}^2 - N\bar{g}_{v1}^2$

$= 2N\theta_{v1}f_{v1}(1 + \theta_{v1}f_{v1}) - N(2\theta_{v1}f_{v1})^2$

$= 2N\theta_{v1}f_{v1}(1 - \theta_{v1}f_{v1})$

If $v \leq S$:

$(G_1'G_2)_{vv} = \underset{\sim}{g}_{v1}'\underset{\sim}{g}_{v2} = -2Nf_{v1}f_{v2}\theta_{v1}\theta_{v2}$

If $v > S$:

$(G_1'G_2)_{vv} = \underset{\sim}{g}_{v1}'\underset{\sim}{g}_{v2} = 2Nf_{v1}f_{v2}(\theta_{v12} - \theta_{v1}\theta_{v2})$

Now, to sum the diagonal elements:

$tr(\text{Cov}[\beta_1, \beta_1]G_1'G_1) = \sum_{v=1}^{V} \frac{h_1^2}{V}\tau_{v1}^2 2N\theta_{v1}f_{v1}(1 - \theta_{v1}f_{v1}) = \frac{Nh_1^2}{V}\sum_{v=1}^{V}\frac{\theta_{v1}(1-\theta_{v1}f_{v1})}{1-f_{v1}}$

$tr(\text{Cov}[\beta_2, \beta_2]G_2'G_2) = \sum_{v=1}^{V} \frac{h_2^2}{V}\tau_{v2}^2 2N\theta_{v2}f_{v2}(1 - \theta_{v2}f_{v2}) = \frac{Nh_2^2}{V}\sum_{v=1}^{V}\frac{\theta_{v2}(1-\theta_{v2}f_{v2})}{1-f_{v2}}$

$tr(\text{Cov}[\beta_1, \beta_2]G_1'G_2) = \sum_{v=1}^{S} \frac{\rho}{V}\sqrt{h_1^2h_2^2}\tau_{v1}\tau_{v2}(-2Nf_{v1}f_{v2}\theta_{v1}\theta_{v2}) = \frac{-N\rho\sqrt{h_1^2h_2^2}}{V}\sum_{v=1}^{S}\frac{\theta_{v1}\theta_{v2}\sqrt{f_{v1}f_{v2}}}{\sqrt{(1-f_{v1})(1-f_{v2})}}$

Thus, we have for heritability in admixed subjects:

$h_{adm}^2 = \text{Var}(G_1\beta_1 + G_2\beta_2) = \frac{h_1^2}{V}\sum_{v=1}^{V}\frac{\theta_{v1}(1-\theta_{v1}f_{v1})}{1-f_{v1}} + \frac{h_2^2}{V}\sum_{v=1}^{V}\frac{\theta_{v2}(1-\theta_{v2}f_{v2})}{1-f_{v2}} - \frac{2\rho\sqrt{h_1^2h_2^2}}{V}\sum_{v=1}^{S}\frac{\theta_{v1}\theta_{v2}\sqrt{f_{v1}f_{v2}}}{\sqrt{(1-f_{v1})(1-f_{v2})}}$
